# Brain dynamics and temporal trajectories during task and naturalistic processing

**DOI:** 10.1101/380402

**Authors:** Manasij Venkatesh, Joseph Jaja, Luiz Pessoa

**Affiliations:** Department of Electrical and Computer Engineering, University of Maryland, College Park, MD, USA; Department of Psychology and Maryland Neuroimaging Center, University of Maryland, College Park MD, USA

## Abstract

Human functional Magnetic Resonance Imaging (fMRI) data are acquired while participants engage in diverse perceptual, motor, cognitive, and emotional tasks. Although data are acquired temporally, they are most often treated in a quasi-static manner. Yet, a fuller understanding of the mechanisms that support mental functions necessitates the characterization of dynamic properties. Here, we describe an approach employing a class of recurrent neural networks called reservoir computing, and show the feasibility and potential of using it for the analysis of temporal properties of brain data. We show that reservoirs can be used effectively both for condition classification and for characterizing lower-dimensional “trajectories” of temporal data. Classification accuracy was approximately 90% for short clips of “social interactions” and around 70% for clips extracted from movie segments. Data representations with 12 or fewer dimensions (from an original space with over 300) attained classification accuracy within 5% of the full data. We hypothesize that such low-dimensional trajectories may provide “signatures” that can be associated with tasks and/or mental states. The approach was applied across participants (that is, training in one set of participants, and testing in a separate group), showing that representations generalized well to unseen participants. Taken together, we believe the present approach provides a promising framework to characterize dynamic fMRI information during both tasks and naturalistic conditions.

## 1 Introduction

Functional Magnetic Resonance Imaging (fMRI) data are acquired while participants engage in diverse perceptual, motor, cognitive, and emotional tasks. Three-dimensional images are acquired every 1-2 seconds and reflect the state of blood oxygenation in the brain, which serves as a proxy for neuronal activation. Although data are acquired temporally, they are most often treated in a quasi-static manner [Huettel et al., 2004]. In blocked designs, fairly constant mental states are maintained for 15-30 seconds, and data are essentially averaged across multiple repetitions of a given block type, such as performing a working memory task. In event-related designs, short trials typically 1-5 seconds long are employed and the responses evoked are estimated with multiple regression.

Many fMRI studies also are constrained spatially, in the sense that activation is analyzed independently at every location in space. However, so-called multivariate pattern analysis techniques capitalize on information that is potentially distributed across space to characterize and classify brain activation [Haxby et al., 2001, Kamitani and Tong, 2005, Haynes and Rees, 2006]. For example, in an early study,Cox and Savoy [Cox and Savoy, 2003] investigated the performance of a linear discriminant classifier, a polynomial classifier, and a linear support vector machine to classify objects presented to participants from voxel activations (i.e., features) across visual cortex. Since then the field has matured and developed a wealth of approaches, including the investigation of “representational” content carried by brain signals [Kriegeskorte et al., 2006]. However, given the relatively low signal-to-noise ratio of fMRI data (which necessitates a large number of repetitions of data segments of interest), the vast majority of multivariate methods for investigating brain data are “static,” that is, the inputs to classification are patterns of activation that are averaged across time (“snapshots”) [Haynes, 2015]. Some studies have proposed using temporal information as well as spatial data [Mourao-Miranda et al., 2007, Hutchinson et al., 2009, Nestor et al., 2011, Janoos et al., 2011, Chu et al., 2011]. One of the goals in such cases has been to extend the features provided for classification by considering a temporal data segment instead of, for example, the average signal during the acquisition period of interest. Despite some progress, key issues remain largely unexplored, including understanding the integration of temporal information across time, and questions about the dimensionality of temporal information (see below).

In all, despite the potential of fMRI to be used to investigate temporal structure in task data the technique is employed in a largely static fashion. However, a fuller understanding of the mechanisms that support mental functions necessitates the characterization of dynamic properties. Here, we describe an approach that aims to address this gap. At the outset, we acknowledge that the low-pass nature of the blood-oxygenation response is such that dynamics should be understood at a commensurate temporal scale (on the order of a few seconds or typically longer). Yet, many mental processes unfold at such time scales, such as the processing of event boundaries [Zacks et al., 2001], a gradually approaching threatening stimulus [Najafi et al., 2017], listening to a narrative [Ferstl et al., 2005], or watching a movie [Hasson et al., 2004].

Several machine learning techniques exist that are sensitive to temporal information. Among them, recurrent neural networks (RNNs) have attracted considerable attention [Williams and Zipser, 1989, Pearlmutter, 1989,Horne and Giles, 1995]. However, effectively training RNNs is very challenging, particularly without large amounts of data ([Pascanu et al., 2013]; but for recent developments see [Martens and Sutskever, 2011, Graves et al., 2013]). Here, we propose to use *reservoir computing* to study temporal properties of fMRI data. This class of algorithms, which includes liquid-state machines [Maass et al., 2002], echo-state networks [Jaeger, 2001, Jaeger and Haas, 2004], and related formalisms [Steil, 2004, Sussillo and Abbott, 2009], includes recurrence (like RNNs) but the learning component is only present in the read-out, or output, layer (Fig. 1A). Because of the feedback connections in the reservoir, the architecture has memory properties, that is, its state depends on the current input and past reservoir states. The read-out stage can be one of many simple classifiers, including linear discrimination or logistic regression, thus providing considerable flexibility to the framework. Intuitively, reservoir computing is capable of separating complex stimuli because the reservoir “projects” the input into a higher-dimensional space, making it easier to classify them. Of course, this is related to the well-known difficulty of attaining separability in low dimensions, as was recognized early on with the use of Perceptrons [Cover, 1965].

**Fig. 1:**
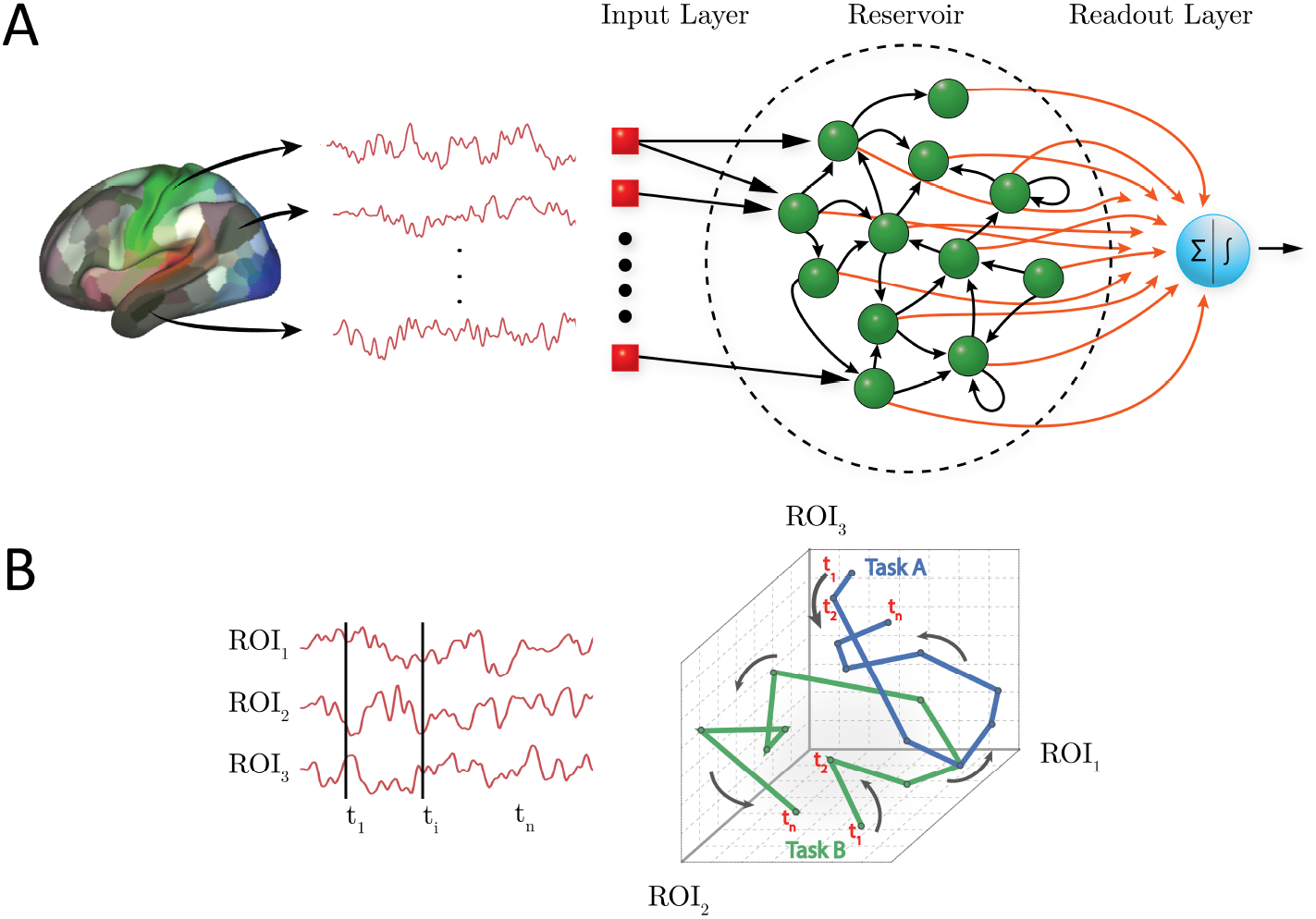
Reservoir computing and temporal trajectories. (A) Brain data are provided to a three-layer neural network. The input layer registers the input; in the present case, activation at time *t* across a set of regions of interest (ROIs). The reservoir layer contains units with random connections, and provides a memory mechanism such that activation at time *t* is influenced by past time points. The readout (output) layer indicates the category of the input; in the present case, the binary labels “0” or “1” corresponding to task/condition. Only the connections between the reservoir and the readout layer (shown in red) are adaptable. (B) Time series data can be represented as a temporal “trajectory.” In the case of data from three ROIs (left), activation can be plotted along axes “1,” “2,” and“3” at each time point *t* (right). In this manner, activation during a hypothetical task exhibits a particular trajectory, whereas activation during a second task exhibits a different trajectory (blue and green lines for Task A and B, respectively). Note that the trajectories might overlap at several time points (the activation at those time points is the same for both tasks), but the entire trajectory provides a potentially unique “signature” for the task/condition in question.

Reservoir computing has been effectively used for time series prediction [Lu et al., 2017], temporal signal classification [Skowronski and Harris, 2007], as well as applications in several other domains [Triefenbach et al., 2010, Vandoorne et al., 2008]. Here, we show the feasibility and potential of using it for the analysis of temporal properties of brain data. The central objectives of our study were as follows. First, to investigate reservoir computing for the purposes of *classifying* fMRI data, in particular when temporal structure might be relevant, including both task data and data acquired during movie watching. The latter illustrates the potential of the technique for the analysis of naturalistic conditions, which are an increasing focus of research. Here, classification was attempted on task condition (for example, theory of mind versus random motion) or movie category (“scary” versus “funny”).

Our second goal was to characterize the *dimensionality* of the temporal information useful for classification. Many systems can be characterized by a lower-dimensional description that captures many important system properties. In neuroscience, research with multi-unit neuronal data has suggested that low-dimensional “*trajectories*” can be extracted from high-dimensional noisy data [Yu et al., 2009, Buonomano and Maass, 2009]. As Yu and colleagues proposed [Yu et al., 2009], a neural trajectory potentially provides a compact representation of the high-dimensional recorded activity as it evolves over time, thereby facilitating data visualization and the study of neural dynamics under different experimental conditions (see also [Gao et al., 2017]). Here, we hypothesized that reservoir computing could be used to extract low-dimensional fMRI trajectories that would provide “signatures” for task conditions and/or states (Fig. 1B). For both of our objectives, we sought to investigate them at the between-participant level (in contrast to within-participant) to ascertain the generalizability of the representations created by the proposed framework.

## 2 Methods

### 2.1 Human Connectome Project Data

We employed working memory and theory of mind data collected as part of the Human Connectome Project (HCP; [Barch et al., 2013]). Data were collected with a TR of 720 ms. We employed data from *N* = 200 participants. This included *N* = 100 unrelated participants, and a separate, non-overlapping set of *N* = 100 participants randomly selected from the *N* = 1200 data release.

#### Working memory dataset

Participants performed a “2-back” working memory task, where they indicated if the current stimulus matched the one presented two stimuli before, or a control condition called “0-back” (without a memory component). Data for two runs were available, each containing four 27.5-second blocks of each kind. Stimuli consisted of faces, places, tools, and body parts. To account for the cue response at the start of the block and the hemodynamic lag, data from 12-30 seconds after block onset were used (25 data points per block).

#### Theory of mind dataset

Participants performed a theory of mind task, where they indicated whether short video clips displayed a potential social interaction, no meaningful interaction (“random”), or they were unsure. Stimuli consisted of 20-second video clips in which geometric objects (squares, circles, triangles) appeared to interact either meaningfully, or randomly. Data for two runs were available, each containing five video clips; thus, five social interaction and five random clips were available in total. To account for hemodynamic lag (no cue was employed), data from 3-21 seconds after block onset were used (25 data points per block).

### 2.2 Participants (movie watching)

Sixteen participants with normal or corrected-to-normal vision and no reported neurological or psychiatric disease were recruited from the University of Maryland community. Data from 12 participants (5 males and 7 females, ages 18-22 years; mean: 20.6, SD: 1.5) were employed for data analysis (two participants voluntarily quit the study before completion, and data from three participants were discarded due to head motion exceeding 4 mm). The project was approved by the University of Maryland College Park Institutional Review Board and all participants provided written informed consent before participation.

### 2.3 Movie data acquisition

Functional and structural MRI data were acquired using a 3T Siemens TRIO scanner with a 32-channel head coil. First, a high-resolution T1-weighted MPRAGE anatomical scan (0.9 mm isotropic) was collected. Subsequently, we collected six functional runs of 384 EPI volumes each using a multiband scanning sequence [Feinberg et al., 2010]. For 3/12 participants, the following imaging parameters were used: TR = 1.25 sec, TE = 42.8 ms, FOV = 210 mm, voxel size: 2.0 mm isotropic, number of slices = 72, and multiband factor = 6. For the remaining 9 participants, slightly altered parameters used were: TR = 1.25 sec, TE = 39.4 ms, FOV = 210 mm, voxel size: 2.2 mm isotropic, number of slices = 66, and multiband factor = 6. For all participants, non-overlapping oblique slices were oriented approximately 20-30 clockwise relative to the AC-PC axis (2.0 mm isotropic) helping to decrease susceptibility artifacts at regions such as the orbitofrontal cortex and amygdala.

### 2.4 Movie data

We employed fMRI data collected from 12 usable participants while viewing short movie segments (duration between 1-3 minutes) with content that was either “scary,” “funny,” or “neutral” (neutral segments were not utilized here) (see Table S1 for a list of the movies employed). Participants viewed one movie clip of each kind per run for a total of six runs. A total of 30 movie clips (15 of each kind) were extracted from the movie segments such that at least one clip originated from each of the movies viewed. Clips contained 25 data points (like the HCP data above), which lasted 31.25 seconds (data were acquired with a TR of 1.25 seconds). All video clips focused on parts of the movie segments that were deemed by one of the authors (M.V.) to be of high arousal/interest.

### 2.5 Preprocessing

#### HCP data

Data were part of the “minimally preprocessed” release, which included fieldmap-based distortion correction, functional to structural alignment, and intensity normalization. Data were collected with a TR of 720 ms. We investigated cortical data which are directly provided in surface representation. In addition to the preprocessing above, we regressed out 12 motion-related variables (6 translation parameters and their derivatives) using the 3dDeconvolve routine of the AFNI package [Cox, 1996] (with the “ortvec” option). Low frequency signal changes were also regressed out with the same routine by using the “polort” option (with the polynomial order set automatically).

#### Movie data

A combination of packages and in-house scripts were used to preprocess both the functional and anatomical MRI data. The first five volumes of each functional run were discarded to account for equilibration effects. Slice-timing correction (with AFNI’s 3dTshift) used Fourier interpolation to align the onset times of every slice in a volume to the first acquisition slice, and then a six-parameter rigid body transformation (with AFNI’s 3dvolreg) corrected head motion within and between runs by spatially registering each volume to the first volume.

To skull strip the T1 high-resolution anatomical image (which was rotated to match the oblique plane of the functional data with AFNI’s 3dWarp), the ROBEX package [Iglesias et al., 2011] was used. Then, FSL’s epi-reg was used to apply boundary-based co-registration in order to align the first EPI volume image with the skull-stripped T1 anatomical image [Greve and Fischl, 2009]. Next, ANTS [Avants et al., 2011] was used to estimate a nonlinear transformation that mapped the skull-stripped anatomical image to the skull-stripped MNI152 template (interpolated to 1-mm isotropic voxels). Finally, ANTS combined the transformations from co-registration (from mapping the first functional EPI volume to the anatomical T1) and normalization (from mapping T1 to the MNI template) into a single transformation that was applied to map volume registered functional volumes to standard space (interpolated to 2-mm isotropic voxels). The overall approach described in this paragraph was based on [Smith et al., 2018] and used previously by our group ([Najafi et al., 2017]). The resulting spatially normalized functional data were smoothed using a 6 mm full-width half-maximum Gaussian filter. Spatial smoothing was restricted to gray-matter mask voxels (with AFNI’s 3dBlurInMask). Finally, the average intensity at each voxel (per run) was scaled to 100.

### 2.6 Regions of Interest

#### HCP data

Because our goal was to evaluate the general framework described here, and not test specific hypotheses tied to particular brain regions, we considered cortical data only. Because for cortical data the HCP processing pipeline is oriented toward a surface representation, we employed the cortical parcellation developed by their research group [Glasser et al., 2016]. The parcellation includes 360 cortical regions of interest (ROIs), and is based on a semi-automated approach that delineates areas based on architecture, function, connectivity, and topography (see Fig. S1A).

#### Movie data

ROIs were determined in a volumetric fashion. To do so, we employed a simple *k*-means clustering algorithm that generated 500 cortical ROIs. Specifically, clustering was based on the {*x, y, z*} spatial coordinates of voxels in cortex (not their time series), and an *L*_2_ distance metric was employed to favor the grouping of nearby voxels (see Fig. S1B). We also performed our analysis with 400 and 600 ROIs and observed essentially the same results; thus, the precise choice of the number of ROIs does not appear to be critical. In addition to the cortical ROIs, given the importance of the amygdala for emotional processing in general, we also included two amygdala ROIs (one per hemisphere). Each ROI was generated by combining the lateral and the central/medial amygala (as defined in [Nacewicz et al., 2012]) into a single region.

For both HCP and movie data, a summary ROI-level time series was obtained by averaging signals within the region.

### 2.7 Reservoir computing

For temporal data analysis, we adopted the reservoir formulation used in echo-state networks [Jaeger, 2001, Jaeger and Haas, 2004]. The general reservoir computing architecture includes three main elements: an input layer, a reservoir, and a read-out (or output) layer (Fig. 1A). The input layer registers the input and is connected with the reservoir. The reservoir contains units that are randomly interconnected within the reservoir, as well as connected to units in the read-out layer.

Only connections to the read-out layer undergo learning. Here, the input layer activations, **u**(*t*), represented activation for the condition of interest at time *t*. The number of input units corresponded to the number of ROIs, and one value was input with every data sample (every time *t*). The output layer contained a single unit with activation corresponding to a category label (0 or 1, coding the task condition). At every time step, the activations of the reservoir units were updated, determining a reservoir state, **x**(*t*), and the readout, *z*(*t*), was instantiated. Thus, the input time series data generated an output time series (one per time point) corresponding to category labels.

The state of the reservoir was determined by [Lukoševičius, 2012]

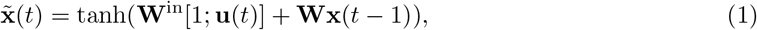

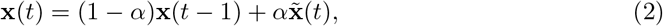

where **x̃**(*t*) is an intermediate state. The function tanh(*x*) was applied element-wise and implemented a sigmoidal activation function. The notation [*·*;*·*] stands for vertical vector concatenation. Both **x̃**(*t*) and 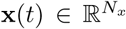 where the dimensionality of the reservoir *N_x_* = *τ × N_u_* is determined by the number of input units, *N_u_*, and the parameter *τ*. The dimensionality of the reservoir, *N_x_*, is related to the memory of the reservoir, namely, the number of past data points that can influence the current output. A general rule of thumb is that for an input of size *N_u_*, to remember *τ* time points in the past, the reservoir should have size at least *τ* × *N_u_* [Lukoševičius, 2012]. The weight matrices 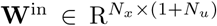 and 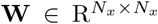 are the input-to-reservoir and within-reservoir matrices, respectively. The parameter *α* ∈ (0, 1] is the leakage (or “forgetting”) rate. Interpreting the equations above, **x̃**(*t*) is a function of a weighted contribution of the input plus a weighted contribution of the prior reservoir state (passed through a sigmoidal function) (Equation 1). The reservoir state, **x**(*t*), is a weighted average of the previous reservoir state **x**(*t –* 1) and ̃(*t*) based on weights (1 – *α*) and *α*, respectively (Equation 2). Overall, this reservoir formulation allows it to encode temporal information in a spatial manner, that is, across the reservoir units. The present reservoir implementation utilized code from the Modeling Intelligent Dynamical Systems research group (http://minds.jacobs-university.de/research/esnresearch/).

A key idea in reservoir computing is that the weight matrices **W**^in^ and **W** are not trained, but instead generated randomly (unlike RNNs which include adaptable weights in all layers). The non-symmetric matrix **W** is typically sparse with nonzero elements obtained from a standard normal distribution, *𝒩*(0, 1); here, of the *N_x_* × *N_x_* matrix entries, 10*N_x_* were randomly chosen to be non-zero. The input matrix **W**^in^ is generated according to the same distribution, but typically is dense. It is crucial to ensure that the largest absolute value of the eigenvalues of the reservoir weight matrix **W** be less than 1, as this ensures the “echo state” property [Jaeger, 2001]: the state of the reservoir, **x**(*t*), should be uniquely defined by the fading history of the input, **u**(*t*).

### 2.8 Classification

The reservoir state, **x**(*t*), can be viewed as a random non-linear high-dimensional expansion of the input signal, **u**(*t*). If the inputs are not linearly separable in the original space, 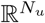, they often become separable in the higher dimensional space, 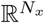, of the reservoir. Such so-called “kernel tricks” are common in machine learning algorithms [Murphy, 2012, Scholkopf and Smola, 2001], and reservoirs embed that property within a temporal processing context.

The read-out layer of a reservoir architecture can employ one of multiple simple components, including linear or logistic regression, or support vector machines. Here, we employed *l*_2_-regularized logistic regression with a constant inverse regularization parameter, *C* = 1 [Pedregosa et al., 2011], for two-class classification. Given a set of data points and category labels, a logistic regression classifier learns the weights of the output layer, **W**^out^, by maximizing the conditional likelihood of the labels given the data. A gradient descent algorithm searches for optimal weights such that the probability *P* [*z*(*<u>t</u>*<u>) = 1</u>*<u>|</u>***<u>x</u>**(*t*)] = *σ*(**W**^out^**x**(*t*)) is large when **x**(*t*) belongs to class “1” and small otherwise; 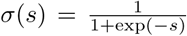 is a logistic function. The classes considered here were “2-back” vs. “0-back” for working memory, “social” vs. “random” for theory of mind, and “scary” vs. “funny” for movie clips.

Because we were interested in temporal properties, classification was performed at every time *t*, with a single classifier. Thus, as stated above, the input time series data generated an output time series corresponding to category labels, **z**(*t*).

Finally, note that our objective was to characterize the capabilities of the reservoir framework to capture temporal information for classification as a function of time. Accordingly, we employed the “minimal” classification machinery at the output end of our algorithm. Had the objective been to maximize classification values, we could have included, for example, a “second classifier” (that is, one after the readout layer) that considered simultaneously all classification values **z**(*t*) during the block, for example.

### 2.9 Dimensionality reduction

Functional MRI data are very high-dimensional if one considers all the voxels or surface coordinates acquired with standard imaging parameters. Typical anatomical parcellations considerably reduce the dimensionality as 100 to 1,000 ROIs are usually employed (and one time series is commonly employed per ROI). Whereas this represents a major reduction in dimensionality, it is important to understand if lower-dimensional characterizations of the data are informative. Here, we sought to determine classification accuracy of temporal fMRI data of lower-dimensional representations. In particular, what is the lowest dimensionality that provides performance comparable to that obtained with the “full” dimensionality? Recall that because reservoir states, **x**(*t*), are non-linear high-dimensional expansions of the input signals, **u**(*t*), their dimensionality is higher than the number of ROIs (by the factor *τ*; see above).

For dimensionality reduction, we employed principal component analysis (PCA) to the reservoir states, **x**(*t*) (Fig. 2A). In brief, PCA provides a coordinate transformation such that the dimensions are orthogonal. In the new coordinate system, the transformed reservoir state, **y**(*t*), has the same dimensionality as the original representation. It is possible to reduce the dimensionality of the input by retaining a subset of the dimensions that capture the most variance of the original signals. Our goal, however, was to perform dimensionality reduction while considering dimensions that were useful for classification, and not necessarily capturing the most variance. To do so, we performed logistic regression analysis using PCA-transformed states, **y**(*t*), and used the weights of the resulting classifier to select the principal components that best distinguished the task conditions (somewhat akin to partial least squares; see [Brereton and Lloyd, 2014]). Components associated with large positive weights encourage the decision toward one of the classes, whereas those associated with large negative weights encourage the decision toward the other class. We can then select the *k* dimensions with largest positive weights and the *k* dimensions with the largest negative weights, which we called the “top” and “bottom” principal components; we called time series data along the *k* dimensions “top and bottom time series.” For example, the dimension with the largest positive weight (call it dimension 1) is associated with time series *y*_1_(*t*). See (Fig. 2A) for a schematic of the sequence of data transformations. Importantly, since these components were determined by using classifier weights, which were based solely on training data, test data were unseen and could be used to assess classification performance (see Section 3.2.).

**Fig. 2:**
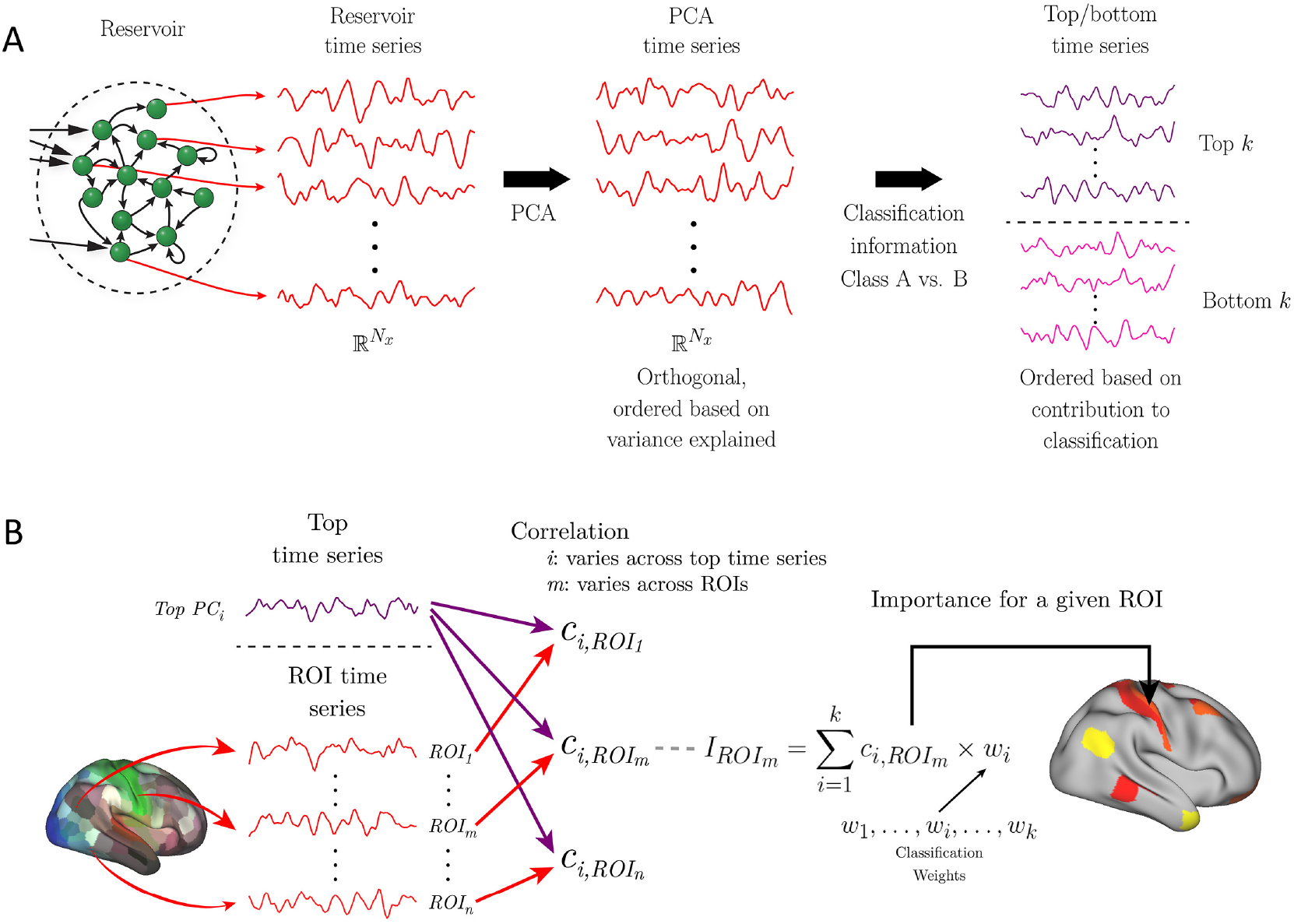
Dimensionality reduction and brain activation. (A) Reservoir activation provides a high-dimensional “expansion” of the input vectors at every time *t*. In this manner, the reservoir is associated with a reservoir time series of dimensionality *N_x_* = *τ × N_u_*, where *τ* is a parameter and *N_u_* is the size of the input vector (here, the number of ROIs employed). The first step of dimensionality reduction employed principal component analysis (PCA). Subsequently, the dimensions were ordered based on the weights of the logistic regression classifier (the larger the absolute value of the weight, the more important the dimension for classification). We refer to the data along these dimensions as “top” (indicative of one output category) and “bottom” (indicative of the alternative category) time series. (B) To indicate brain regions expressing top time series information, each top time series (at left, only one is shown for simplicity) was correlated with the original fMRI time series of each ROI. The correlations along with the weights associated with the top time series are indicative of the *importance* of an ROI to classifying a condition (as the active condition). A set of ROIs can then be indicated on the brain (right) that express the *k* top time series based on the importance values, *I*_ROI_. For example, the *l* ROIs with largest importance values can be shown, or those ROIs such that the importance exceeds a specific threshold. Taken together, although the reservoir time series representation is a high-dimensional expansion of the input data, it is possible to map the brain regions that most express the top time series, which are the ones providing the greatest contribution to classification.

#### Region importance

The high-dimensional representation of the reservoir, or the lower-dimensional representation of the *k* top/bottom components, is considerably removed from the original fMRI time series. However, it is important to determine which original ROI time series express the most information about them, what we call *region importance*. To do so, we first computed the Pearson correlation between each original ROI fMRI time series and each of a number of top time series. To facilitate interpretation of importance, we employed only top time series because they contributed positively to classification performance, that is, they had positive classification weights (Fig. 2A); recall that positive weights provided evidence for the “active class” and negative weights for the control condition.

The contribution of an ROI to classification was not only dependent on its correlation with a top time series but also the logistic regression weight associated with the time series. Specifically, the weight *w_i_* from the PCA-transformed reservoir dimension, *y_i_*(*t*). Thus, the “importance value” of an ROI to a particular task condition was based on the correlation value times the classification weight (Fig. 2B). Finally, an importance index for an ROI was obtained by adding the extent to which an ROI time series “loaded” (correlated with) onto *k* top time series corresponding to the task (*k* was 5 for working memory data, 6 for theory of mind data, and 6 for movie data; see Results for explanation of how *k* was determined). Importance values were then shown on brain maps (for illustration, we display the 25 highest importance values/ROIs on the brain). For display of importance across tasks, values were rescaled into the range [0, 1]: 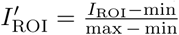, where *I*_ROI_ is the importance value prior to rescaling.

### 2.10 Additional temporal analyses

To understand the ability of reservoirs to integrate information across time, temporal information was also used in a straightforward manner. Here, the activations across a block were concatenated into a single long vector of size **number-of-ROIs** × **number-of-time-points**. The resulting vector was then used as input to a logistic regression classifier (instead of data at each time step separately) and performance determined.

To assess the role of the non-linear expansion in the reservoir, we compared the results with those obtained with a linear autoregressive model, a standard technique used to model time series data. Activations at time *t* for each ROI *k*, *u_k_*(*t*), were predicted based on the previous *p* time points, such that the predicted value at time *t* was given by

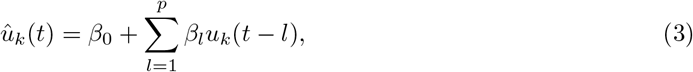
where *p* is the so-called model order. The estimated coefficients, *β_i_*, that minimize the squared error between *u_k_*(*t*) and *û_k_*(*t*) can be obtained via least squares. As routinely done, the first *p* time points in the block were ignored in this AR(*p*) model. The activations predicted based on this model were used to train a logistic classifier, as done with reservoirs.

### 2.11 Statistical approach and tests

#### Studying reservoir parameters

Our initial goal was to investigate the ability of reservoirs to capture temporal information in fMRI data. Accordingly, we varied the parameters *α* (forgetting rate) and *τ* (ratio of the number of reservoir to input units), which together determine the memory properties of the reservoir. To determine classification accuracy, we employed a *between-subject* cross-validation approach. For HCP data, *N* = 100 unrelated participants were used (for reference, we will call this the “first” dataset). Five-fold cross-validation was employed by randomly splitting the data into 80-20 train-validation sets: in each fold, 80 participants were used to train the reservoir, and 20 participants for validation (that is, to determine classification accuracy in unseen data). This procedure was applied for each of the *α* × *τ* parameter combinations.

Because we were interested in temporal properties, classification was performed at every time *t*. Classification accuracy for a block was defined on the time point with the best classification accuracy, *t*_best_, during the block. We did not employ the average accuracy across the entire block, because for temporally varying data some segments of the block would not be expected to contain distinguishing information; for instance, the beginning of a block (see Fig. 6). To improve robustness, we considered *t*_best_ and its two adjacent time points, *t*_best_ − 1 and *t*_best_ + 1, such that accuracy was the average across these three time points. Note that *t*_best_ was defined on training data only and applied on test data that was not used to define it. Overall, the “first” dataset served to investigate reservoir parameters and define the best-performing *α*, *τ*, and *t*_best_.

To evaluate the classification accuracy of reservoirs, we employed permutation testing [Ojala and Garriga, 2010]. Given the computational demands of permutation testing in our framework, *p*-values were based on 1000 iterations (with the exception of the test of randomizing temporal information; see below). The best-performing reservoir parameters were used to train a logistic classifier (see Section‘2.8) by utilizing the entire *N* = 100 participants of the “first” dataset, but accuracy was determined entirely based on a separate *N* = 100 dataset (for reference, the “second” dataset). This ensured that classification information generalized to completely unseen data. The observed accuracy was then compared to a null distribution of accuracy that was obtained by repeating this procedure 1000 times but with class labels randomly permuted; for each iteration, training with permuted labels was performed on the “first” dataset and testing was based on the “second” dataset. If *m* is the number of iterations where the classification accuracy on data with permuted labels exceeded the accuracy on data with true labels, and *k* is the total number of iterations, the *p*-value was obtained as 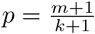

#### Comparison with other methods

We compared the performance observed with reservoirs to three other methods. The first was to simply test classification on raw activation signals. In this case, the logistic classifier was directly fed the inputs **u**(*t*); everything else was identical to the classification with reservoirs. In other words, the inputs to classification were directly from the input layer and not the reservoir (see Fig. 1A). Thus, identical to the case of reservoirs, classification on activation signals generated a time series of corresponding labels *z*(*t*). The other two methods employed temporal information as outlined previously: concatenating data across time points in a block, and using autoregressive modeling. The reservoir used the best-performing *α*, *τ*, and *t*_best_ obtained using the “first” dataset. Likewise, the order of the autoregressive model (*p* = 10) was the best performing one obtained with the “first” dataset; the model orders investigated were *p* = {2, 5, 10}, which were comparable to the reservoir parameter *τ* values, but for results as a function of *p*, see Fig. S2). The actual comparison between methods was established based on the “second” dataset. To compare accuracy values, a Wilcoxon signed-rank test was utilized.

#### Randomizing temporal information

To test whether the temporal order within a block is informative, data points within a block were randomly shuffled. For fMRI data, simply reshuffling breaks the serial dependency in the data, and so a “wavestrapping” approach was used [Bullmore et al., 2004]. In this manner, the autocorrelation structure is preserved by shuffling the wavelet coefficients at each level (which are whitened and therefore exchangeable). Given the computational demands of the procedure, the associated permutation testing was based on 100 iterations.

#### Movie data

For movie data, we only had a limited amount of data. Accordingly, all classification accuracy results were based on 6-fold cross-validation by randomly splitting the data into 10-2 train-validation sets (10 participants for training, 2 participants for testing).

## 3 Results

Initially, we employed Human Connectome Project (HCP) data from two tasks: working memory and theory of mind. Working memory was chosen to represent a task with a relatively stable “cognitive set” (at the time scale of fMRI). For this case, the active condition comprised 25-second blocks of the so-called 2-back memory task, where participants were asked to indicate if the current item matched the one before the immediately preceding one. We employed the 0-back condition as a comparison condition (no working memory requirement). In contrast to working memory, theory of mind data were expected to exhibit some form of dynamics. During the active condition, participants watched 20-second clips containing simple geometrical objects (including squares, rectangles, triangles, and circles) that engaged in a socially relevant interaction (for example, they appeared to initially fight and then make up) that unfolded throughout the duration of the clip. When watching such clips, one has the impression that the potential meaning of the interactions gradually becomes clearer. The baseline condition in this case consisted of same-duration clips of the same geometrical objects following random motion.

To investigate the ability of the reservoir to capture temporal information in fMRI data, we varied the parameters *α* (forgetting rate) and *τ* (ratio of the number of reservoir to input units), which together determine the memory properties of the reservoir. Accuracy in classifying theory of mind task increased as the size of the reservoir increased, and exceeded 90% (Fig. 3B), which robustly differed from chance (permutation test, *p <* 10^−3^). In contrast, for the working memory task, accuracy differed from chance (permutation test, *p <* 10^−3^) but remained essentially the same, showing that enhanced performance was not always simply due to an increase in reservoir size (Fig. 3A).

**Fig. 3:**
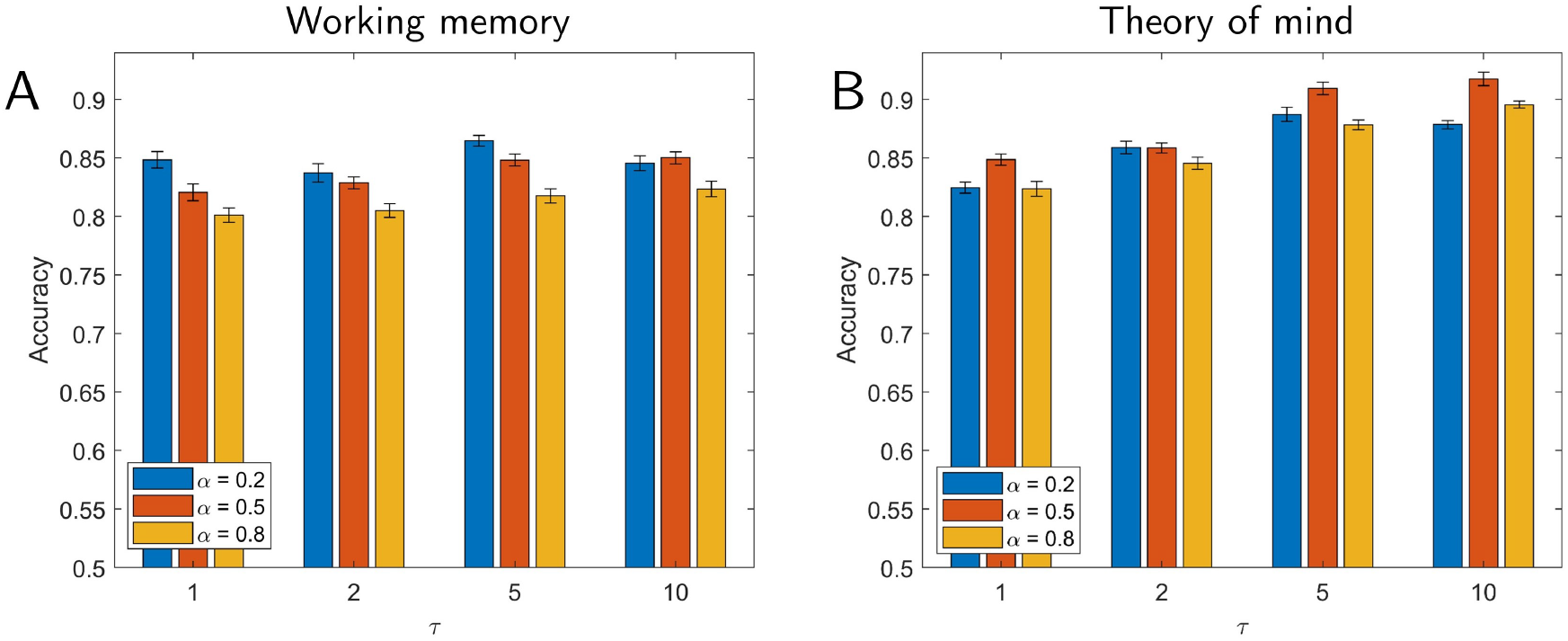
Classification accuracy for working memory (A) and theory of mind (B) tasks. Parameters varied included *α* (which determines the forgetting rate) and *τ* (which determines reservoir size), both of which influence the memory properties of the reservoir. Performance did not vary substantially as a function of reservoir memory for the working memory task (A) but improved as memory increased for the theory of mind task (B). Error bars show the standard error of the mean across validation folds.

We reasoned that if temporal information and context are important, classification should be affected by temporal order, especially in the case of theory of mind data. To evaluate this claim, we trained the classifier without temporal information, namely, by randomly shuffling the data points in a block prior to training (while preserving autocorrelation structure; see Methods); testing was performed with blocks that were temporally ordered. In this case, mean classification accuracy was drastically reduced to 67.1% correct (using the same best-performing parameters) compared to 91.9% correct (permutation test, *p <* 0.01). This further indicates that it was not simply the high-dimensional expansion of the reservoir but also its memory that helped improve classification. For completeness, we also tested classification of working memory data in the same manner. In this case, mean classification accuracy was 74.4%, which was a relatively small (but robust; permutation test, *p <* 0.01) decline in performance relative to the best mean accuracy of 86.3% on the unshuffled working memory data.

### 3.1 Comparisons with other approaches

To better characterize the classification performance of reservoirs, we performed a series of comparisons with simpler schemes. All results in the present section were obtained by evaluating the “second” dataset and are summarized in Table 1. Classification accuracy using raw activation data (no reservoir, that is, **u**(*t*) signals) was 77.6% for working memory and 84.2% for theory of mind. For theory of mind, when the reservoir size was small (*τ* = {1, 2}), accuracy was comparable to that with raw activation (see Fig. 3B). It appears that when the number of reservoir units is relatively small, the reservoir representation of the data is poor, particularly when the forgetting rate is high, possibly due to the inability of the reservoir to generate a satisfactory representation of dynamically changing data with fewer dimensions.

**Table 1:** Classification accuracy for reservoirs and additional processing approaches. The *p*-values were determined via Wilcoxon signed-rank tests comparing classification accuracy of reservoirs to each method.

| Working memory: “2-back” vs. “0-back” |  |  |  |  |  |
| --- | --- | --- | --- | --- | --- |
| | Accuracy | | Accuracy | $p$-value | Signed-rank |
| Reservoirs | 86.3% | Raw activation | 77.6% | $< 10^{-8}$ | 821 |
| | | Concatenation | 82.8% | $< 10^{-3}$ | 761 |
| | | Autoregressive model | 81.1% | $< 10^{-3}$ | 1432 |
| Theory of mind: “social” vs. “random” |  |  |  |  |  |
| | Accuracy | | Accuracy | $p$-value | Signed-rank |
| Reservoirs | 91.9% | Raw activation | 84.2% | $< 10^{-8}$ | 2062 |
| | | Concatenation | 86.6% | $< 10^{-3}$ | 1370.5 |
| | | Autoregressive model | 87.8% | $< 10^{-3}$ | 1172 |
| Movie data: “scary” vs. “funny” |  |  |  |  |  |
| | Accuracy | | Accuracy | $p$-value | Signed-rank |
| Reservoirs | 70.9% | Raw activation | 60.2% | $< 10^{-3}$ | 77 |
|  |  | Concatenation | 65.3% | 0.0151 | 69.5 |
|  |  | Autoregressive model | 64.6% | 0.0107 | 70 |

The next two approaches explicitly considered temporal properties of the data. First, we concatenated activation signals from multiple time steps, and performed logistic classification on the concatenated data. Classification accuracy of working memory data was 82.8% correct and of theory of mind data was 86.6% correct. Next, we sought classification with an autoregressive model, which yielded accuracy of 81.1% correct (working memory) and 87.8% correct (theory of mind) (both of which were obtained with a model of order *p* = 10). Although performance with these two methods was relatively close to that with reservoirs, the latter was consistently superior (see Table 1). Finally, note that the classification values across methods used in the present study were rather stable, as illustrated by the comparison of the estimates based on cross-validation (“first” dataset) and those of the “second” dataset (Table S2); recall that all statistical results were based entirely on the “second” dataset which was never used for parameter selection.

### 3.2 Low-dimensional representation

We sought to investigate the dimensionality of the reservoir representation capable of classifying fMRI data. To do so, we performed PCA on reservoir data and determined the number of principal components required to achieve classification performance similar to that on the full data. Instead of considering components in terms of the variance explained, we considered “top” and “bottom” components based on how they improved classification (see Methods; Fig. 2A). Fig. 4 shows classification accuracy as the number of components was increased from 2 to 20 in steps of 2 (one top and one bottom component were added together at a time). For working memory, 10 principal components (5 top and 5 bottom) were required to attain classification at 95% of the level of the full dimensionality; for theory of mind, 12 principal components (6 top and 6 bottom) were required. Note that these components captured only 7% and 8% of the total variance of the working memory and theory of mind datasets, respectively, which should be compared to 72% and 71% captured by first 10 and 12 components when they were selected based on the amount of variance explained (and not classification), consistent with the idea that a relatively small percentage of the original signal variance was informative for classification.

**Fig. 4:**
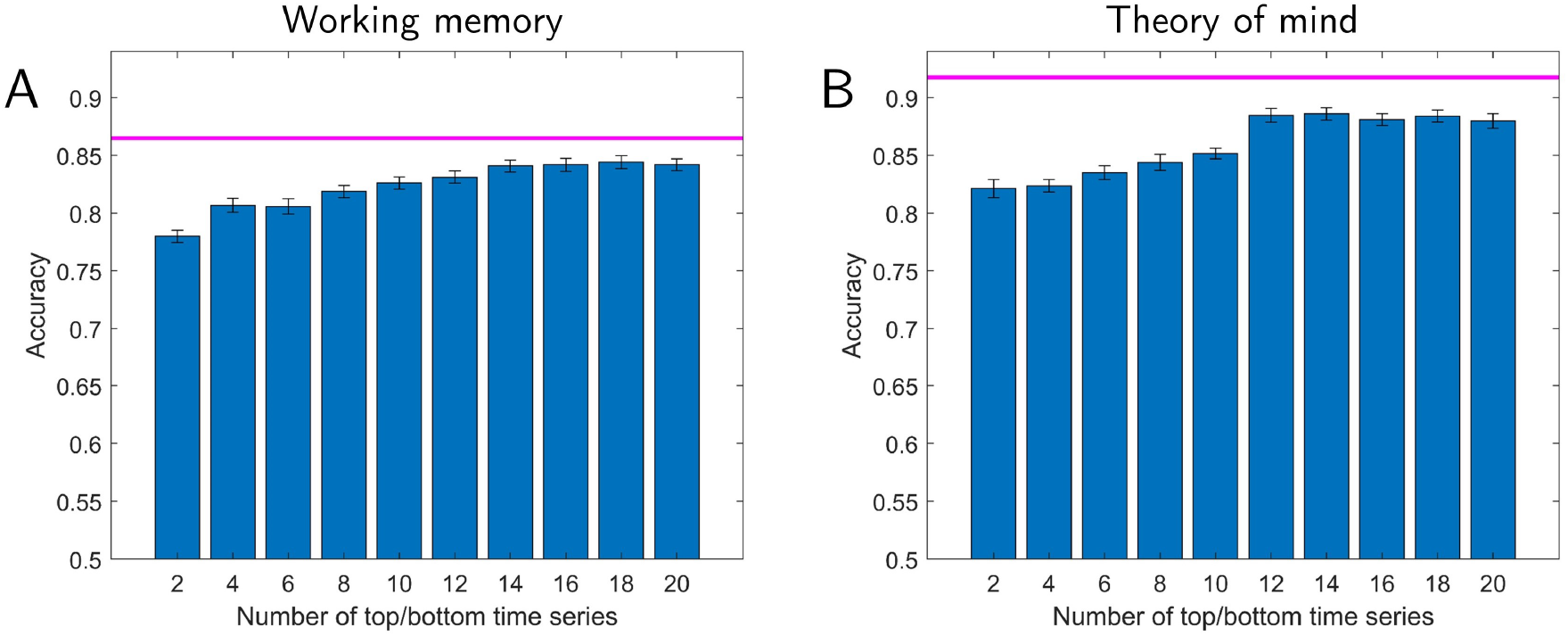
Lower-dimensional representation of reservoir signals and classification accuracy. Accuracy is shown as a function of the number of top plus bottom components. The magenta line indicates the highest performance using all components. Classification accuracy with a lower-dimensional representation reached within 95% of the the full data with 10 and 12 dimensions for working memory (A) and theory of mind (B), respectively. Error bars show the standard error of the mean across validation folds.

Fig. 4 also shows that classification with only the top/bottom 2-4 components attained accuracy at approximately 90% of that obtained with the full dimensionality. We could thus capitalize on this property and select three components so as to visualize their trajectories as a function of time (Fig. 5). For working memory data, the trajectories indicated that the two conditions should exhibit better-than-chance classification even at the beginning of the block. In contrast, for theory of mind data, the trajectories of the social and random conditions initially overlapped, but later became quite distinct. To qualify these observations, we plotted classification accuracy as a function of time during task blocks (for various reservoir configurations). Fig. 6 shows the results for the full dimensionality; results for the top/bottom components are displayed in Fig. S3. For working memory, accuracy was initially around 70% correct, and increased gradually up to 85% for the best reservoir configuration. For theory of mind, accuracy was initially at chance, and increased more abruptly between time points 5-8 (3.5-5.5 seconds), eventually attaining classification over 90%.

**Fig. 5:**
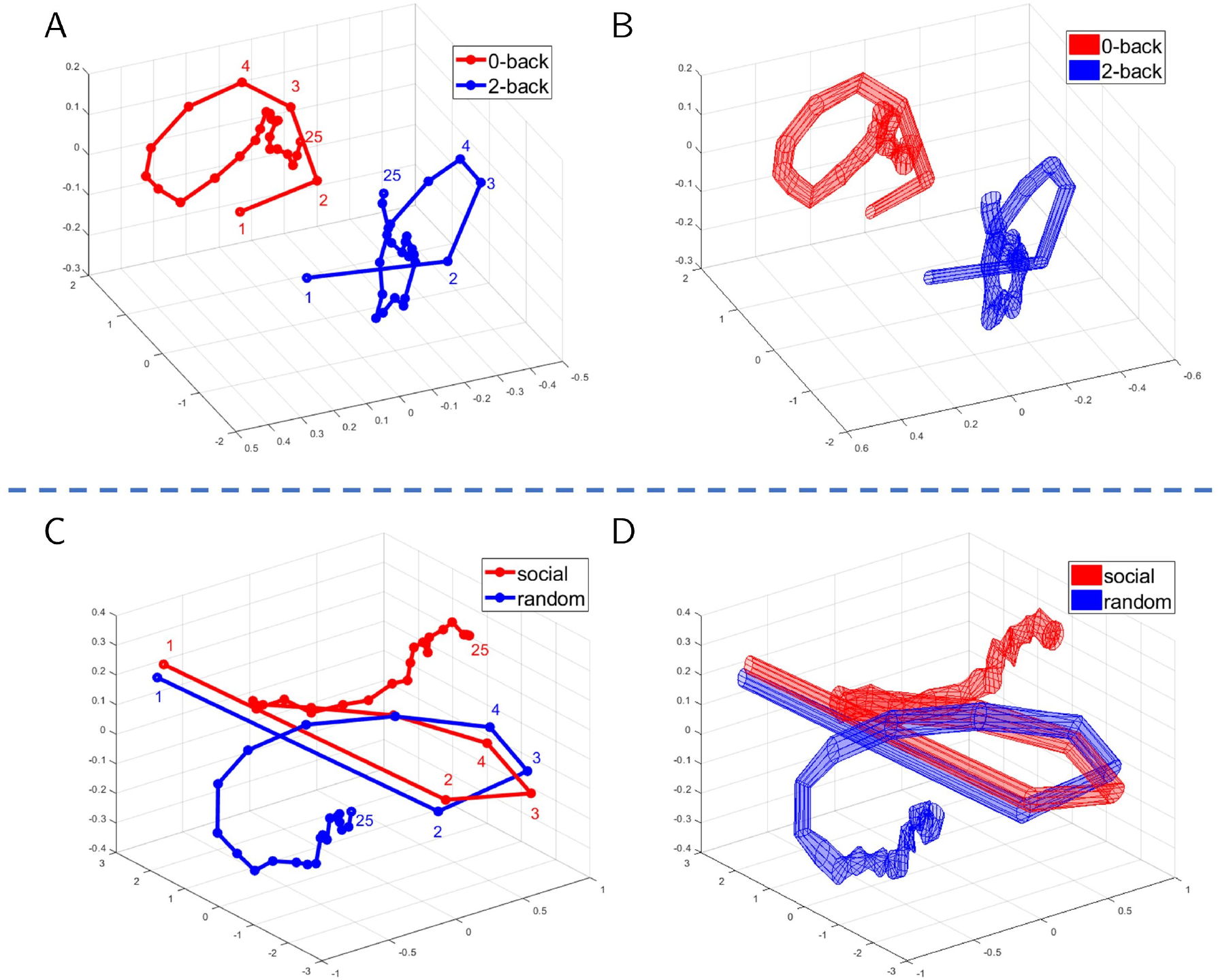
Temporal trajectories for task fMRI data. Mean trajectories are displayed in (A) for working memory and (C) for theory of mind. Variability (standard error across participants) is displayed in (B) and (D), respectively. For working memory data (A-B), the trajectories were well separated throughout the block. For theory of mind data (C-D), the trajectories initially overlapped but diverged after 6-7 points. Trajectories were based on the top three principal components.

**Fig. 6:**
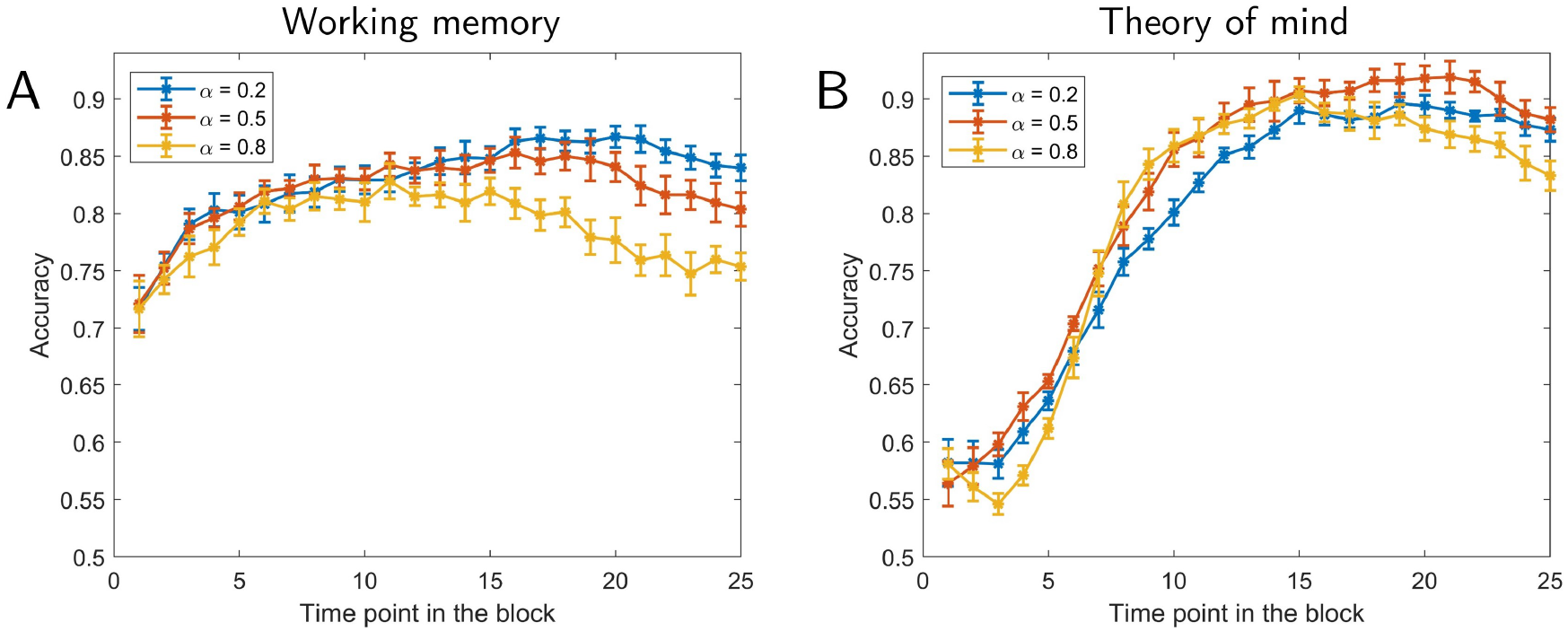
Classification accuracy as a function of time. Results for working memory (A) and theory of mind (B). Accuracy is shown as a function of time point within a task block. Different curves show results for different forgetting rates, *α*. The values of *τ* were based on the parameters exhibiting highest accuracy in Fig. 3. Error bars show the standard error of the mean across validation folds.

How does classification performance based on a lower-dimensionality representation compare to that obtained with regions previously reported to be engaged by theory of mind? To investigate this issue, we used ROIs from a meta-analysis of prior fMRI studies [Schurz et al., 2014], and selected those found to be engaged during social animation tasks. The results based on the 22 ROIs from the meta-analysis are shown in Fig. 7. Performance mostly leveled off with *τ* = 5 at around 80% correct. Note that this performance was lower than that observed with the top/bottom 10 dimensions by about 5%. It is also instructive to compare the results obtained with the meta-analysis ROIs to those with the full data, with the latter exhibiting classification accuracy about 10% higher. The results with the the meta-analysis ROIs did not change appreciably even when the size of the reservoir was increased to match the much larger reservoir size used with the full dimensionality (this was the case when *τ* = 165; recall that the size of the reservoir is given by *τ* times the size of the input vector). Thus, inferior performance with meta-analysis ROIs was not simply due to the size of the reservoir.

**Fig. 7:**
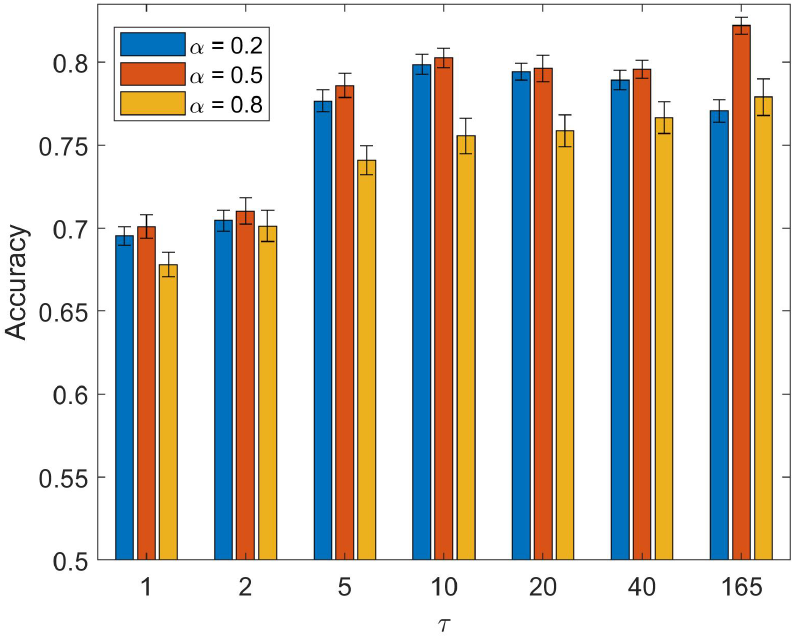
Classification accuracy for theory of mind data with regions involved in “social animation” (based on a meta-analysis of fMRI studies). Performance largely leveled off with a parameter *τ* = 5 or larger. Because the meta-analysis only included 22 regions of interest, we increased *τ* so as to match the reservoir size with that used with the full dimensionality (*τ* = 165). Note that accuracy was limited to around 80% even when the reservoir was large, indicating that the limiting factor was not the size of the reservoir. Error bars show the standard error of the mean across validation folds.

Finally, we also investigated the low-dimensional representation obtained using principal components directly based on activation signals (Fig. 8). For working memory, a small number of components (3 top and 3 bottom) attained classification at 95% of the level of full activation data. However, for theory of mind, 28 components (14 top and 14 bottom) were required. This was more than twice of what was required for the reservoir data indicating that they captured more information required for classification in fewer dimensions. For completeness, Fig. S4 shows temporal trajectories when principal components were based on activation data; it appears that these do not provide temporal signatures as informative as with reservoirs.

**Fig. 8:**
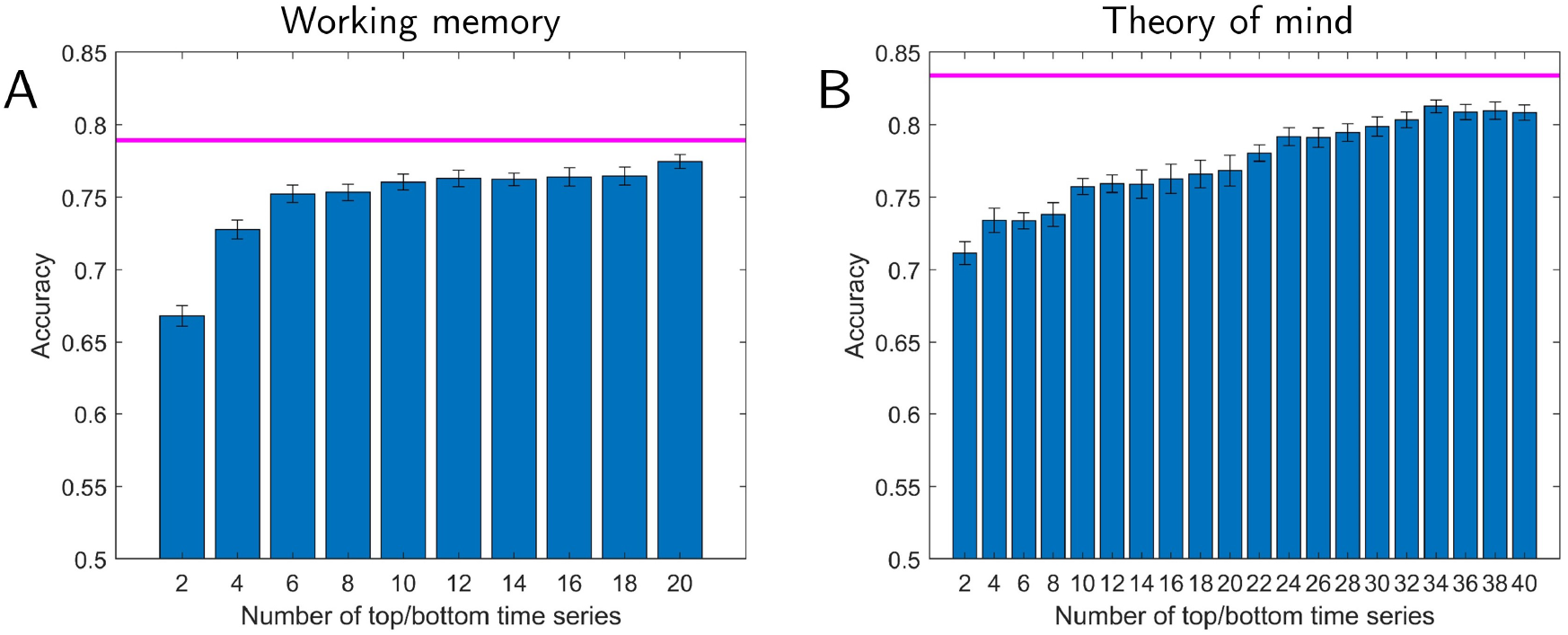
Lower-dimensional representation of activation data (no reservoir) and classification accuracy. Accuracy is shown as a function of the number of top plus bottom components. The magenta line indicates the highest performance using the full-dimensional activation data. For working memory (A), classification accuracy with the lower-dimensional representation reaches 95% of the full data with 6 dimensions. However, for the theory of mind (B), 28 dimensions (as opposed to 12 when using reservoir data) of the lower-dimensional representation. Error bars show the standard error of the mean across validation folds.

### 3.3 Mapping low-dimensional representations to the brain

We sought to determine the brain regions providing the greatest contributions to classification (see also [Shine et al., 2018]). To do so, we computed an importance index for each ROI based on time series data (see Fig. 2B). Fig. 9 illustrates some of the ROIs supporting classification for the working memory and theory of mind tasks selected based on the highest importance values. For this analysis, we used the top 5 time series for working memory and top 6 for theory of mind (the top components that were part of the those attaining 95% classification accuracy relative to the full dimensionality, as discussed in the previous section).

**Fig. 9:**
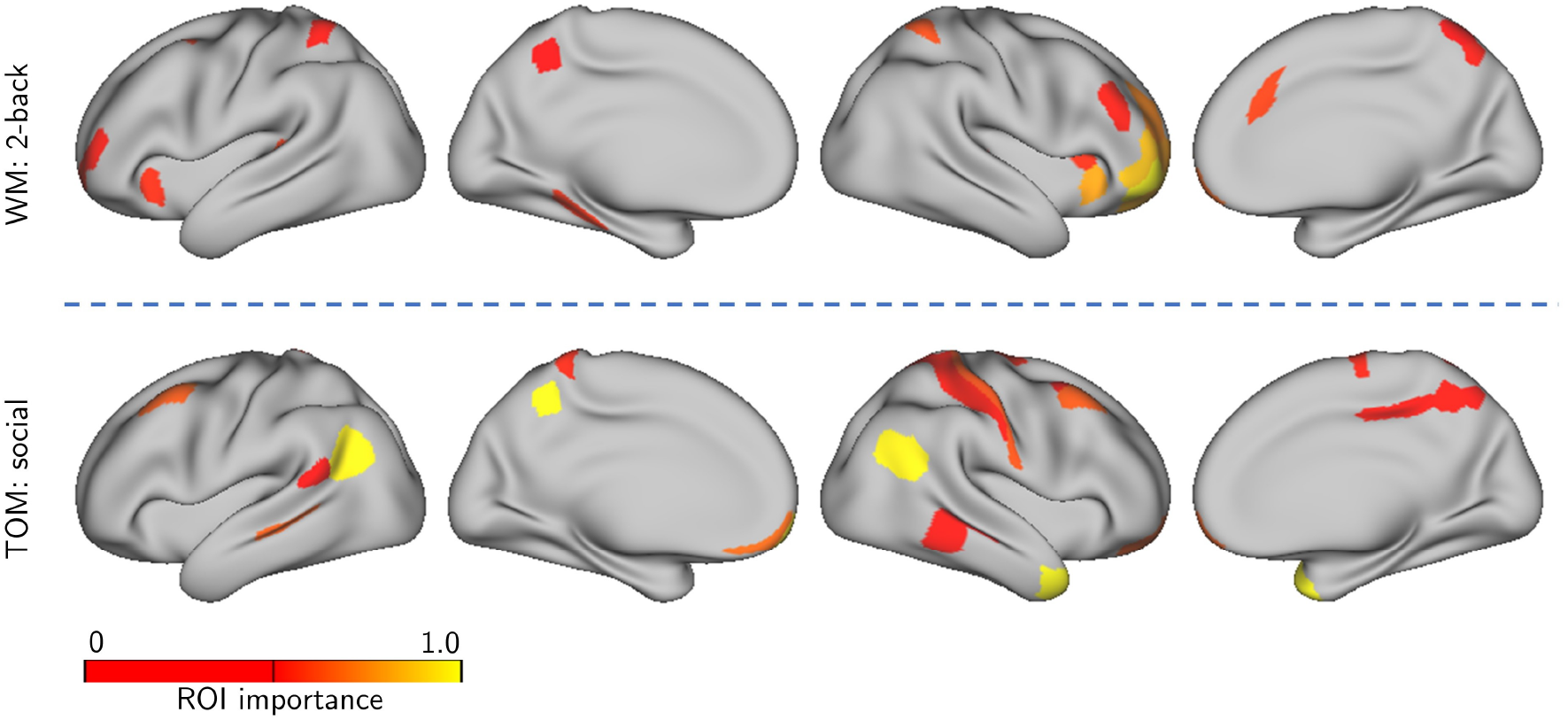
Importance maps for task data. Lower-dimensional time series representation expressed on the brain. The colored ROIs are those with original fMRI time series expressing (“loading on”) “top” time series the most (see Fig. 2 for details).(A) Regions supporting “2-back” in the classification of working memory data. (B) Regions supporting “social” in the classification of theory of mind data.

### 3.4 Movie clips

We further investigated our framework by attempting to classify data segments extracted from movies (31.25 seconds long). Twelve usable participants viewed short movie clips (between 1-3 minutes long; see Methods) of scary or funny content. Given the emotional content of the clips, we added left and right amygdala ROIs to the set of cortical ones. Classification accuracy (“scary” vs. “funny” clips) is displayed in Fig. 10A and reached around 70% correct for larger reservoirs (which was robustly above chance levels; permutation test, *p <* 10^−3^). Like in the case of theory of mind data, performance improved with larger reservoirs. The accuracy for individual movies was between 60% and 80%, showing that classifier performance was not driven by one or a few of the movies watched. In addition, we compared classification with reservoirs with that obtained with activation signals (no reservoir; 60.2%), concatenated data (65.3%), and an autoregressive model (64.6%). As in the case of task data, reservoirs performed best, although the numerical difference was relatively modest (Table 1).

**Fig. 10:**
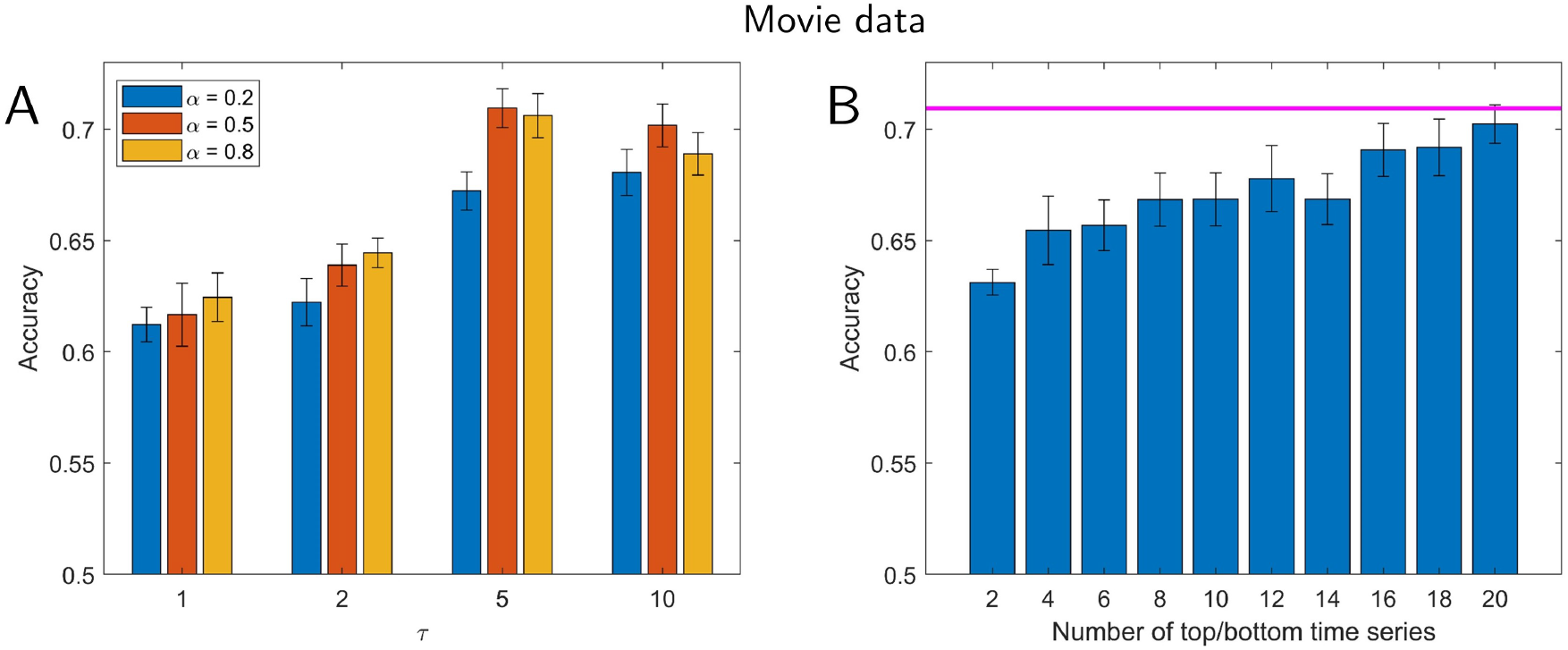
Classification for movie clips. Participants viewed short movie clips that were either scary or funny. (A) Accuracy as a function of the parameters *α* (which determines the forgetting rate) and *τ* (which determines reservoir size). (B) Lower-dimensional representation and classification accuracy. Accuracy is shown as a function of the number of “top” plus “bottom” components. The magenta line indicates the highest performance using all components. Classification accuracy with a lower-dimensional representation reached that of the full data with around 20 dimensions, and reached within 95% of the the full data with 12 dimensions. Error bars show the standard error of the mean across validation folds.

We also investigated lower-dimensional representations of movie data (Fig. 10B). Classification accuracy with 20 dimensions (out of 502) performed at the same level as with the full dimensionality, and with 12 dimensions within 95% of that with all dimensions. In a more exploratory fashion, we investigated temporal trajectories during movie watching. We compared trajectories generated from individual scary clips with the average trajectory observed for funny movie segments. Some scary clips exhibited trajectories that diverged from the mean trajectory for funny clips earlier on, whereas some diverged later in time (Fig. 11), properties that were also apparent in the time course of classification accuracy values. To determine brain regions that most contributed to classification, we computed the importance index for each ROI as with task data. Fig. 12 illustrates some of the brain regions involved when we used the top 6 time series, which attained 95% classification accuracy relative to the full dimensionality (see Fig. 10).

**Fig. 11:**
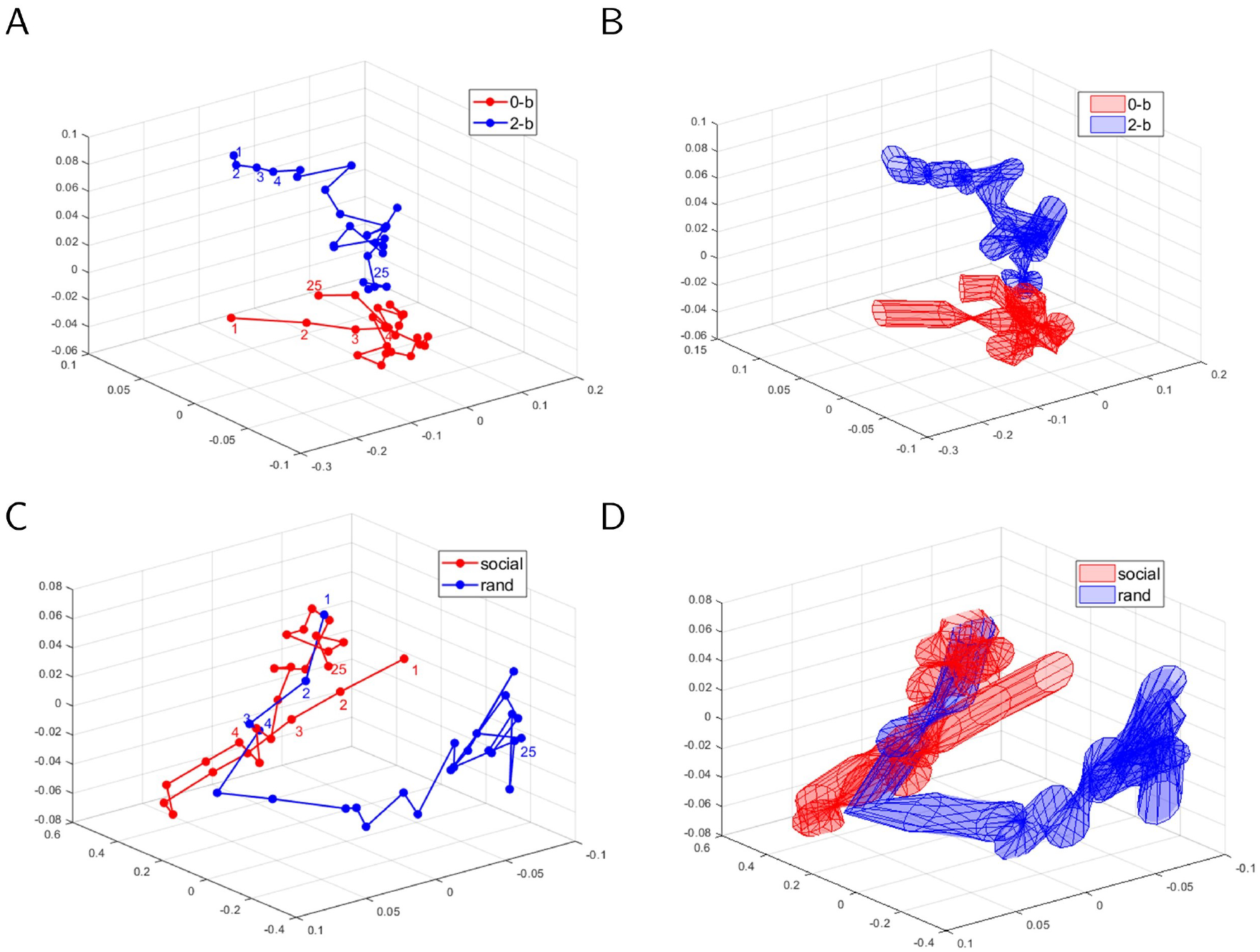
Temporal trajectories for sample clips in the movie data. The red trajectories are for particular “scary” movie clips whereas the blue trajectories are averaged across all “funny clips”. Mean trajectories are displayed in (A) and (C) for two particular scary movie clips. Variability (standard error across participants) is displayed in (B) and (D), respectively. For the movie clip in (A), the trajectories started to separate later than they do for the movie clip in (C). An analysis of the accuracy of these clips as a function of time revealed similar properties. Trajectories were based on the “top” three principal components.

**Fig. 12:**
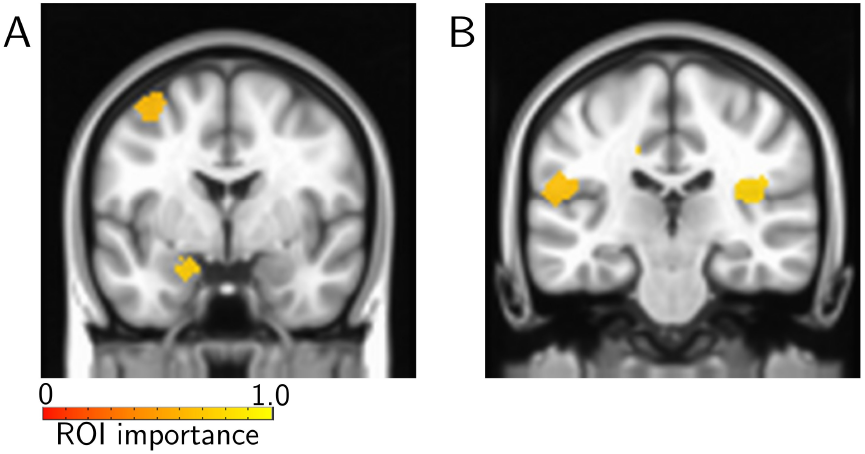
Importance maps for movie data. Lower-dimensional time series representation expressed on the brain. The colored ROIs are those with original fMRI time series expressing (“loading on”) “top” time series the most (see Fig. 2 for details). Regions supporting “scary” included the left amygdala (A) and the insula (B).

## 4 Discussion

In the present paper, we sought to analyze fMRI data with reservoir computing which, like recurrent neural networks, is a technique developed to process temporal data. We show that reservoirs can be used effectively for temporal fMRI data, both for classification and for characterizing lower-dimensional trajectories of temporal data. Importantly, the approach was performed in an out-of-sample fashion, namely, performance was only evaluated in participants whose data were not included for training, demonstrating that the representations of reservoirs generalized well across participants.

### 4.1 Investigating temporal structure of brain data

To date, most analyses of fMRI brain data focus on understanding relatively static information (but see Introduction for further discussion). Neuronal data acquired with physiology is also most often analyzed in terms of averaged responses during certain response epochs that are believed to be behaviorally relevant. Yet, brain processes are highly dynamic and current understanding would benefit from frameworks that focus on understanding temporal processing (see [Buonomano and Maass, 2009]). Here, we employed reservoir computing to investigate and characterize temporal information in fMRI data. But the framework is sufficiently general that it can be employed with other types of brain data time series, including those from cell electrophysiology, calcium imaging, EEG, and MEG (for the use of reservoirs in other neuroscience applications, see for example [Enel et al., 2016]).

We selected two tasks from the HCP dataset to evaluate the model. The working memory task was selected as it was thought to not have a noteworthy temporal component; in the context of classification, the working memory condition was presumed to involve a relatively stable“cognitive set” (in fact, during scanning participants were informed which condition they were performing at the beginning of a block). In contrast, the theory of mind condition was expected to rely more on temporal integration of information (for every trial, participants actively attempted to discern the meaning of the vignettes so as to classify stimuli between meaningful and random).

Although classification of working memory data was a little better than obtained with activation signals alone, classification did not improve with the size of the reservoir, consistent with the notion that temporal information during the block did not play a notable role in performance. In contrast, classification of data segments with meaningful social interactions (vs. random; theory of mind data) benefited from increased reservoir size. With larger memory size, accuracy improved close to 10% in some cases. As stated, the social interactions displayed in the clips build up after a few seconds and evolve throughout the block (for example, two objects “invite” a third to participate in an activity, and all three engage in it). Neuroimaging studies of theory of mind and social cognition have employed such dynamic stimuli to probe the brain correlates supporting this type of processing (for a review, see [Schurz et al., 2014]). Classification between social and random clips was initially at chance levels, and increased sharply within the first 5 seconds of the clip. In future applications of the approach described here, it would be valuable to investigate how individual-level classification performance is potentially associated with behavioral performance and individual differences in social-cognitive skills (see [Bartz et al., 2011]).

Whereas increases in reservoir size did not benefit working memory classification, theory of mind classification improved for theory of mind data, consistent with integration of information across time being useful for classification. Larger reservoir sizes allow signals at the same time *t* to interact in richer ways, too (for example, higher-dimensional signal interactions are possible). Therefore, it is possible that processing theory of mind benefited from this aspect as well (more so than working memory data), and that the correlates of theory of mind data are more distributed in the brain, and of higher inherent dimensionality (see below).

To help understand the behavior of reservoirs for classification of fMRI data, we compared the method to other temporal schemes. Although classification based on both concatenated time series data and autoregressive models performed well for working memory and theory of mind tasks, performance with reservoirs was superior to both approaches. It should be said, however, that quantitatively the improvement was relatively modest. Nevertheless, our results suggest that the non-linear expansion of the reservoir, in addition to its temporal properties, contribute to classification performance. It should be stressed that reservoirs are straightforward to train, unlike other recurrent neural networks with fully adaptable weights. Finally, our general framework also suggests that reservoir computing provides an effective methodology to study lower-dimensional representations of the data, which may provide useful dynamic “signatures” of temporal information of fMRI data (see below).

We also investigated our proposal with naturalistic stimuli, specifically, short clips obtained from movies with either scary or funny content. Classification accuracy increased with larger reservoirs, consistent with the notion that temporal information was useful for distinguishing between the two types of clip. In the context of fMRI data which originate from hemodynamic processes with relatively slow dynamics, we suggest that the reservoir framework developed here might be particularly useful in characterizing temporal processing of naturalistic stimuli, including movies and narratives [Hasson et al., 2004, Lerner et al., 2011].

### 4.2 Low-dimensional trajectories

Brain data collected with multiple techniques, including cell-activity recordings and fMRI, are often of high dimensionality. For example, calcium imaging records neuronal activation across hundreds of neurons simultaneously (for example, [Barbera et al., 2016]). In fMRI, signals from tens or even hundreds of thousands of spatial locations are acquired if whole-brain imaging is considered. Even in the case where only a set of regions is of central interest, hundreds of spatial locations may be involved. Therefore, understanding lower-dimensional representations of signals is important. An important working hypothesis in cell data is that low-dimensional neural trajectories provide compact descriptions of underlying processes [Yu et al., 2009, Buonomano and Maass, 2009].

Here, we investigated lower-dimensionality representations of reservoir states by determining classification accuracy as a function of the number of dimensions employed. For both working memory and theory of mind data, considerable reduction was attained and 12 or fewer dimensions were needed to attain classification at 95% of that obtained with the full data. Furthermore, as illustrated in Fig. 5, even maintaining only three dimensions captured important characteristics of the ability to distinguish task conditions. More generally, we hypothesize that such low-dimensional trajectories may provide “signatures” that can be associated with tasks and/or mental states. We propose that investigating how trajectories differ across different groups of individuals (for example, low vs. high anxiety, autism vs. typically developing, etc.) is a fruitful avenue for future research. Notably, the low-dimensional trajectories captured important temporal properties of the data. For example, for theory of mind data, trajectories were very close initially and diverged subsequently, paralleling the increase from lower to higher classification levels. These results are consistent with the idea that reservoirs provide a mechanism for the accumulation of information over time, and hence result in better accuracy in the later periods of the block.

We investigated how the dimensions with the highest contributions to distinguishing conditions were expressed in the brain by generating importance maps. In the case of working memory, several regions in lateral prefrontal cortex, parietal cortex, and anterior insula contributed to classification. These results are consistent with a large literature showing the participation of these regions in effortful cognitive functions, including working memory [Corbetta and Shulman, 2002, Pessoa and Ungerleider, 2004]. In the case of the theory of mind task, we observed regions in the vicinity of the temporal-parietal junction and associated regions that have been implicated in theory of mind more generally, and the interpretation of social animations in particular [Schurz et al., 2014]. Of interest, regions in the cuneus/pre-cuneous, which are engaged in theory of mind tasks [Schurz et al., 2014, Carrington and Bailey, 2009], were observed, too. Together, these results show that the framework developed here captures information from brain regions known to participate in the tasks investigated.

For the theory of mind data, we further compared classification accuracy obtained with the whole brain ROI partition (360 ROIs) and the lower-dimensional representations, separately, with those obtained by selecting regions from a meta-analysis across studies using social animations [Schurz et al., 2014]. Intriguingly, classification with 22 targeted ROIs performed around 10% lower than obtained with the full data; it also performed more poorly than a lower-dimensional representation with only the top/bottom 4 time series. These results raise the intriguing possibility that regions not detected in the meta-analysis carry useful information about the type of theory of mind investigated here. Therefore, to the extent that classification accuracy relies on features that are “representational,” these results indicate that the correlates of theory of mind are more distributed across the brain. However, given that the present work did not determine the precise features contributing to classification, further work is needed to establish this possibility (see [Haynes, 2015] for discussion of related issues). At the same time, we should note that lower-dimensional representations performed rather well in classifying the stimuli; therefore, representations based on a relatively low number of dimensions (for example, around 10) are feasible. For a related approach to understand the dimensionality of temporal representations in the brain, see [Shine et al., 2018].

We also studied lower-dimensional representations and temporal trajectories obtained from naturalistic movie watching. Whereas this component of our work was more exploratory, our findings revealed that the framework proposed here has the potential to be useful in these scenarios. We not only found that lower-dimensional representations could capture most of the information required for classification, but that temporal trajectories were also informative. Future work should evaluate more systematically the use of our proposal when heterogeneous stimulus sets are employed, such as the movie data investigated here.

In summary, in the present paper, we developed an approach employing reservoir computing, a type of recurrent neural network, and show the feasibility and potential of using it for the analysis of temporal properties of brain data. The framework was applied to both Human Connectome Data and data acquired while participants viewed naturalistic movie segments. We show that reservoirs can be used effectively for temporal fMRI data, both for classification and for characterizing lower-dimensional “trajectories” of temporal data. Importantly, robust classification was performed across participants (in contrast to within-participant classification). We hypothesize that low-dimensional trajectories may provide “signatures” that can be associated with tasks and/or mental states. Taken together, the present approach may provide a flexible and powerful framework to characterize dynamic fMRI information, which can be readily applied to other types of brain data.

## Acknowledgements

M.V. is supported by a fellowship by the Brain and Behavior Initiative, University of Maryland, College Park. L.P. is supported by the National Institute of Mental Health (R01 MH071589 and R01 MH112517). We thank Dustin Moraczewski for discussions about regions of interest linked to theory of mind, Srikanth Padmala for feedback on the manuscript, and Christian Meyer and Anastasiia Khibovska for assistance with figures. Task data were provided by the Human Connectome Project, WU-Minn Consortium (Principal Investigators: David Van Essen and Kamil Ugurbil; 1U54MH091657) funded by the 16 NIH Institutes and Centers that support the NIH Blueprint for Neuroscience Research; and by the McDonnell Center for Systems Neuroscience at Washington University.

## Supplemental figures

**Table S1:**
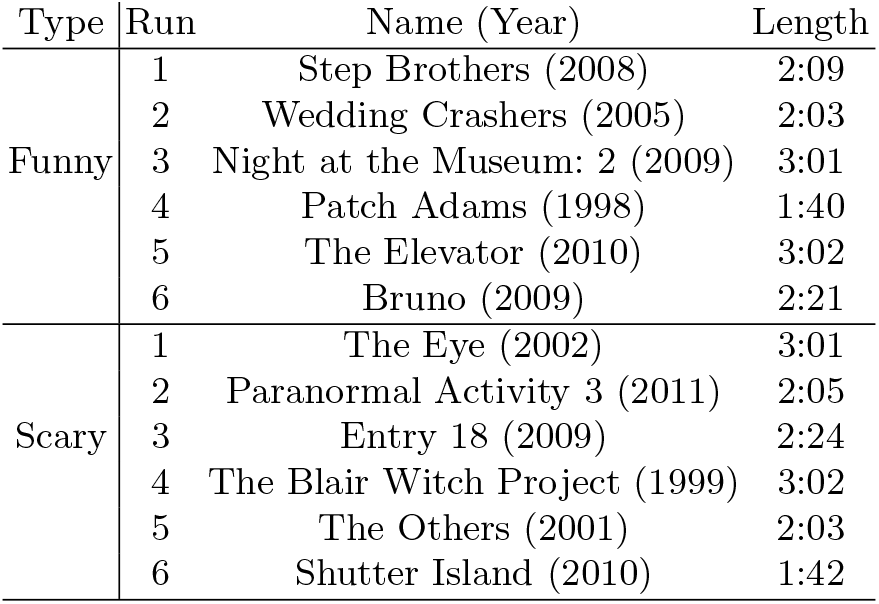
Film names and clip duration for the “scary” and “funny” conditions.

| Type | Run | Name (Year) | Length |
| --- | --- | --- | --- |
| Funny | 1 | Step Brothers (2008) | 2:09 |
|  | 2 | Wedding Crashers (2005) | 2:03 |
|  | 3 | Night at the Museum: 2 (2009) | 3:01 |
|  | 4 | Patch Adams (1998) | 1:40 |
|  | 5 | The Elevator (2010) | 3:02 |
|  | 6 | Bruno (2009) | 2:21 |
| Scary | 1 | The Eye (2002) | 3:01 |
|  | 2 | Paranormal Activity 3 (2011) | 2:05 |
|  | 3 | Entry 18 (2009) | 2:24 |
|  | 4 | The Blair Witch Project (1999) | 3:02 |
|  | 5 | The Others (2001) | 2:03 |
|  | 6 | Shutter Island (2010) | 1:42 |

**Fig. S1:**
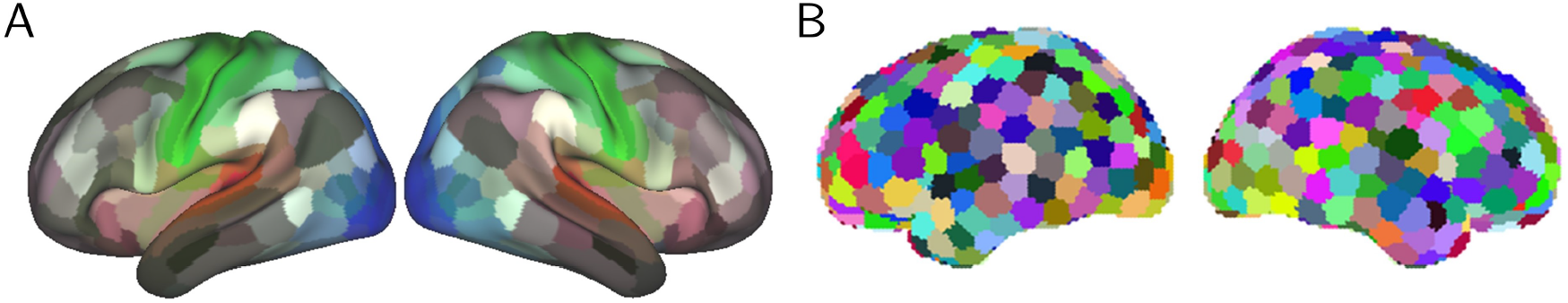
Region of interest (ROI) masks. (A) For Human Connectome Project data, 360 cortical ROIs as provided by [Glasser et al., 2016] were used. (B) For movie data, 500 cortical ROIs obtained from using *k*-means clustering on the spatial coordinates {*x, y, z*} of cortical voxels were used. Two amygdala regions (one per hemisphere) were also included but are not show here.

**Table S2:** Comparison of mean cross-validation accuracy using the “first” dataset and classification accuracy on the “second” dataset. The similar results on the two datasets indicate the robustness of the cross-validation scheme.

| Working memory: “2-back” vs. “0-back” |  |  |
| --- | --- | --- |
|  | Accuracy on the “first” set | Accuracy on the “second” set |
| Reservoirs | 86.5% | 86.3% |
| Raw activation | 78.9% | 77.6% |
| Concatenation | 82.0% | 82.8% |
| Auto-regressive model | 81.2% | 81.1% |
| Theory of mind: “social” vs. “random” |  |  |
|  | Accuracy on the “first” set | Accuracy on the “second” set |
| Reservoirs | 91.8% | 91.9% |
| Raw activation | 83.4% | 84.2% |
| Concatenation | 85.9% | 87.8% |
| Auto-regressive model | 87.2% | 86.6% |

**Fig. S2:**
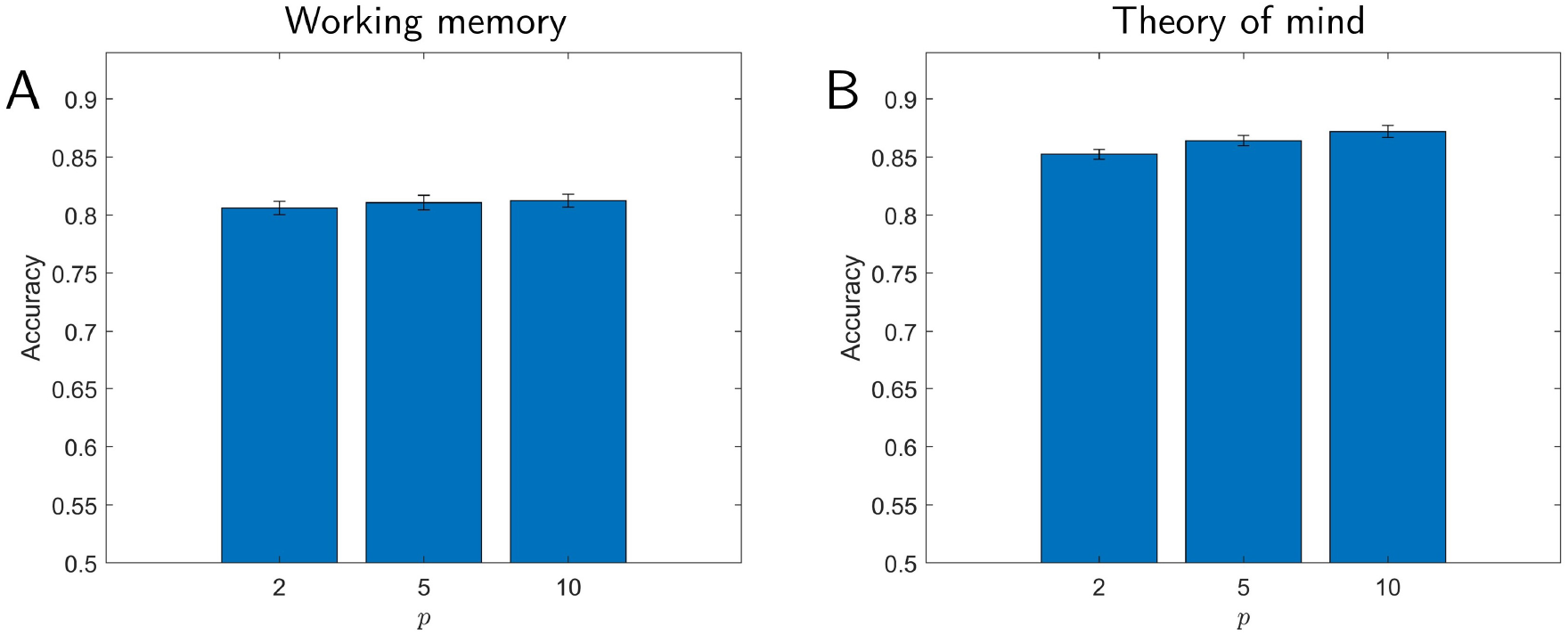
Classification accuracy using autoregressive models for working memory (A) and theory of mind (B). Results are shown as a function of model order, *p*.

**Fig. S3:**
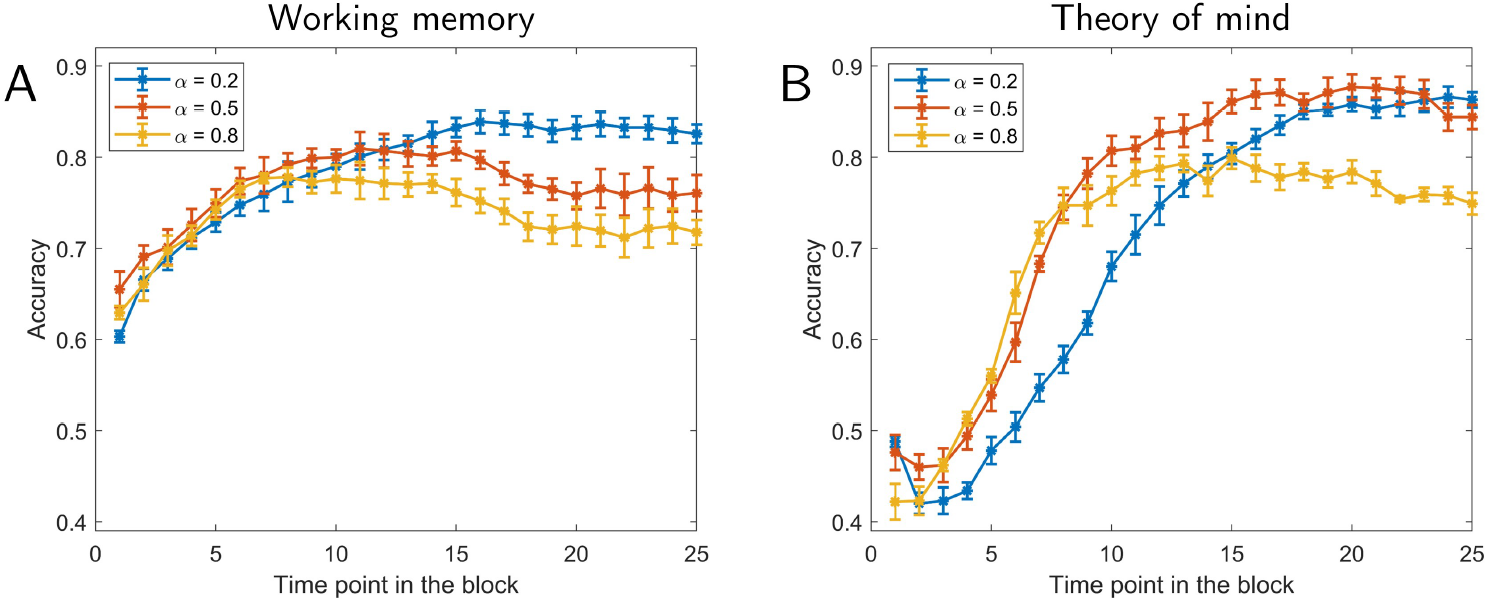
Classification accuracy as a function of time using only the low-dimensional data (10 top/bottom principal components for working memory, and 12 top/bottom principal components for theory of mind). Results for working memory (A) and theory of mind (B). Accuracy is shown as a function of time point within a task block. Different curves show results for different forgetting rates, *α*. The values of *τ* were based on the parameters exhibiting highest accuracy in Fig. 3. Error bars show the standard error of the mean.

**Fig. S4:**
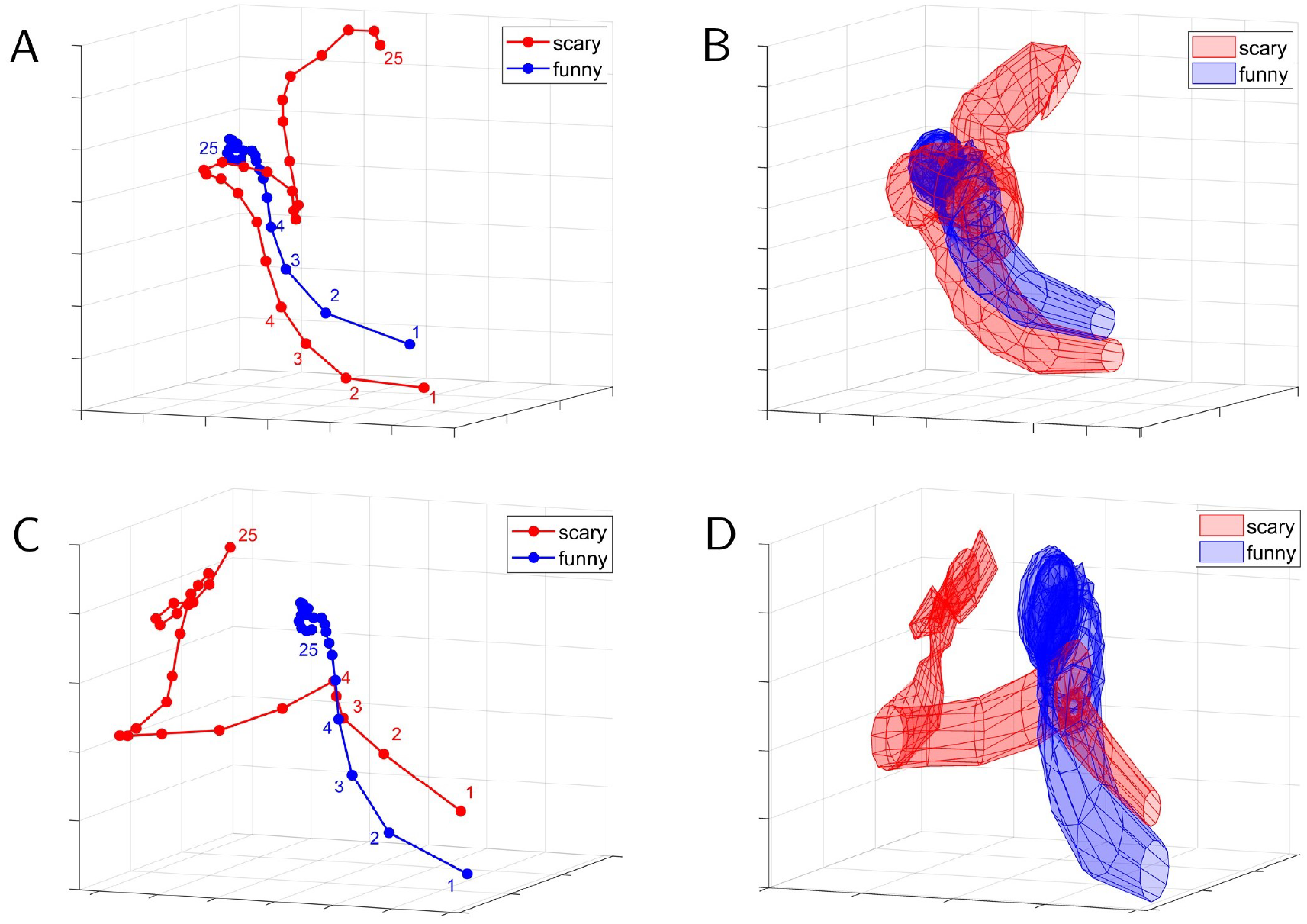
Temporal trajectories based on the “top” three principal components of the input time series (that is, no reservoir) for fMRI task data. Mean trajectories are displayed in (A) for working memory and (C) for theory of mind. Variability (standard error across participants) is displayed in (B) and (D), respectively. For working memory data (A-B), these trajectories were well separated throughout the block. However, for theory of mind data (C-D), the trajectory for the social condition did not evolve temporally as seen when using reservoirs; note that the final states (see point 25) were close to the initial ones (see points 1-2). Therefore, it appears that a low-dimensional representation based directly on input activations does not adequately capture the temporal evolution structure associated with the social condition.

## References

Avants, B. B., Tustison, N. J., Song, G., Cook, P. A., Klein, A., and Gee, J. C. (2011). A reproducible evaluation of ants similarity metric performance in brain image registration. Neuroimage, 54(3):2033–2044.

Barbera, G., Liang, B., Zhang, L., Gerfen, C. R., Culurciello, E., Chen, R., Li, Y., and Lin, D.-T. (2016). Spatially compact neural clusters in the dorsal striatum encode locomotion relevant information. Neuron, 92(1):202–213.

Barch, D. M., Burgess, G. C., Harms, M. P., Petersen, S. E., Schlaggar, B. L., Corbetta, M., Glasser, M. F., Curtiss, S., Dixit, S., Feldt, C., et al. (2013). Function in the human connectome: task-fmri and individual differences in behavior. Neuroimage, 80:169–189.

Bartz, J. A., Zaki, J., Bolger, N., and Ochsner, K. N. (2011). Social effects of oxytocin in humans: context and person matter. Trends in cognitive sciences, 15(7):301–309.

Brereton, R. G. and Lloyd, G. R. (2014). Partial least squares discriminant analysis: taking the magic away. Journal of Chemometrics, 28(4):213–225.

Bullmore, E., Fadili, J., Maxim, V., Sendur, L., Whitcher, B., Suckling, J., Brammer, M., and Breakspear, M. (2004). Wavelets and functional magnetic resonance imaging of the human brain. Neuroimage, 23:S234–S249.

Buonomano, D. V. and Maass, W. (2009). State-dependent computations: spatiotemporal processing in cortical networks. Nature Reviews Neuroscience, 10(2):113.

Carrington, S. J. and Bailey, A. J. (2009). Are there theory of mind regions in the brain? a review of the neuroimaging literature. Human brain mapping, 30(8):2313–2335.

Chu, C., Mourãao-Miranda, J., Chiu, Y.-C., Kriegeskorte, N., Tan, G., and Ashburner, J. (2011). Utilizing temporal information in fmri decoding: classifier using kernel regression methods. Neuroimage, 58(2):560–571.

Corbetta, M. and Shulman, G. L. (2002). Control of goal-directed and stimulus-driven attention in the brain. Nature reviews neuroscience, 3(3):201.

Cover, T. M. (1965). Geometrical and statistical properties of systems of linear inequalities with applications in pattern recognition. IEEE transactions on electronic computers, (3):326–334.

Cox, D. D. and Savoy, R. L. (2003). Functional magnetic resonance imaging (fmri)brain reading: detecting and classifying distributed patterns of fmri activity in human visual cortex. Neuroimage, 19(2):261–270.

Cox, R. W. (1996). Afni: software for analysis and visualization of functional magnetic resonance neuroimages. Computers and Biomedical research, 29(3):162–173.

Enel, P., Procyk, E., Quilodran, R., and Dominey, P. F. (2016). Reservoir computing properties of neural dynamics in prefrontal cortex. PLoS computational biology, 12(6):e1004967.

Feinberg, D. A., Moeller, S., Smith, S. M., Auerbach, E., Ramanna, S., Glasser, M. F., Miller, K. L., Ugurbil, K., and Yacoub, E. (2010). Multiplexed echo planar imaging for sub-second whole brain fmri and fast diffusion imaging. PloS one, 5(12):e15710.

Ferstl, E. C., Rinck, M., and Cramon, D. Y. v. (2005). Emotional and temporal aspects of situation model processing during text comprehension: An event-related fmri study. Journal of cognitive Neuroscience, 17(5):724–739.

Gao, P., Trautmann, E., Byron, M. Y., Santhanam, G., Ryu, S., Shenoy, K., and Ganguli, S. (2017). A theory of multineuronal dimensionality, dynamics and measurement. bioRxiv, page 214262.

Glasser, M. F., Coalson, T. S., Robinson, E. C., Hacker, C. D., Harwell, J., Yacoub, E., Ugurbil, K., Andersson, J., Beckmann, C. F., Jenkinson, M., et al. (2016). A multi-modal parcellation of human cerebral cortex. Nature, 536(7615):171–178.

Graves, A., Mohamed, A.-r., and Hinton, G. (2013). Speech recognition with deep recurrent neural networks. In Acoustics, speech and signal processing (icassp), 2013 ieee international conference on, pages 6645–6649. IEEE.

Greve, D. N. and Fischl, B. (2009). Accurate and robust brain image alignment using boundary-based registration. Neuroimage, 48(1):63–72.

Hasson, U., Nir, Y., Levy, I., Fuhrmann, G., and Malach, R. (2004). Intersubject synchronization of cortical activity during natural vision. science, 303(5664):1634–1640.

Haxby, J. V., Gobbini, M. I., Furey, M. L., Ishai, A., Schouten, J. L., and Pietrini, P. (2001). Distributed and overlapping representations of faces and objects in ventral temporal cortex. Science, 293(5539):2425–2430.

Haynes, J.-D. (2015). A primer on pattern-based approaches to fmri: principles, pitfalls, and perspectives. Neuron, 87(2):257–270.

Haynes, J.-D. and Rees, G. (2006). Neuroimaging: decoding mental states from brain activity in humans. Nature Reviews Neuroscience, 7(7):523.

Horne, B. G. and Giles, C. L. (1995). An experimental comparison of recurrent neural networks. In Advances in neural information processing systems, pages 697–704.

Huettel, S. A., Song, A. W., McCarthy, G., et al. (2004). Functional magnetic resonance imaging, volume 1. Sinauer Associates Sunderland.

Hutchinson, R. A., Niculescu, R. S., Keller, T. A., Rustandi, I., and Mitchell, T. M. (2009). Modeling fmri data generated by overlapping cognitive processes with unknown onsets using hidden process models. NeuroImage, 46(1):87–104.

Iglesias, J. E., Liu, C.-Y., Thompson, P. M., and Tu, Z. (2011). Robust brain extraction across datasets and comparison with publicly available methods. IEEE transactions on medical imaging, 30(9):1617–1634.

Jaeger, H. (2001). The echo state approach to analysing and training recurrent neural networks-with an erratum note. Bonn, Germany: German National Research Center for Information Technology GMD Technical Report, 148(34):13.

Jaeger, H. and Haas, H. (2004). Harnessing nonlinearity: Predicting chaotic systems and saving energy in wireless communication. science, 304(5667):78–80.

Janoos, F., Machiraju, R., Singh, S., and Morocz, I. Á. (2011). Spatio-temporal models of mental processes from fmri. Neuroimage, 57(2):362–377.

Kamitani, Y. and Tong, F. (2005). Decoding the visual and subjective contents of the human brain. Nature neuroscience, 8(5):679.

Kriegeskorte, N., Goebel, R., and Bandettini, P. (2006). Information-based functional brain mapping. Proceedings of the National academy of Sciences of the United States of America, 103(10):3863–3868.

Lerner, Y., Honey, C. J., Silbert, L. J., and Hasson, U. (2011). Topographic mapping of a hierarchy of temporal receptive windows using a narrated story. Journal of Neuroscience, 31(8):2906–2915.

Lu, Z., Pathak, J., Hunt, B., Girvan, M., Brockett, R., and Ott, E. (2017). Reservoir observers: Model-free inference of unmeasured variables in chaotic systems. Chaos: An Interdisciplinary Journal of Nonlinear Science, 27(4):041102.

Lukoševičius, M. (2012). A practical guide to applying echo state networks. In Neural networks: Tricks of the trade, pages 659–686. Springer.

Maass, W., Natschläager, T., and Markram, H. (2002). Real-time computing without stable states: A new framework for neural computation based on perturbations. Neural computation, 14(11):2531–2560.

Martens, J. and Sutskever, I. (2011). Learning recurrent neural networks with hessian-free optimization. In Proceedings of the 28th International Conference on Machine Learning (ICML-11), pages 1033–1040. Citeseer.

Mourao-Miranda, J., Friston, K. J., and Brammer, M. (2007). Dynamic discrimination analysis: a spatial–temporal svm. NeuroImage, 36(1):88–99.

Murphy, K. P. (2012). Machine Learning: A Probabilistic Perspective. The MIT Press.

Nacewicz, B. M., Angelos, L., Dalton, K. M., Fischer, R., Anderle, M. J., Alexander, A. L., and Davidson, R. J. (2012). Reliable non-invasive measurement of human neurochemistry using proton spectroscopy with an anatomically defined amygdala-specific voxel. Neuroimage, 59(3):2548–2559.

Najafi, M., Kinnison, J., and Pessoa, L. (2017). Dynamics of intersubject brain networks during anxious anticipation. Frontiers in human neuroscience, 11:552.

Nestor, A., Plaut, D. C., and Behrmann, M. (2011). Unraveling the distributed neural code of facial identity through spatiotemporal pattern analysis. Proceedings of the National Academy of Sciences, 108(24):9998–10003.

Ojala, M. and Garriga, G. C. (2010). Permutation tests for studying classifier performance. Journal of Machine Learning Research, 11(Jun):1833–1863.

Pascanu, R., Mikolov, T., and Bengio, Y. (2013). On the difficulty of training recurrent neural networks. In International Conference on Machine Learning, pages 1310–1318.

Pearlmutter, B. A. (1989). Learning state space trajectories in recurrent neural networks. Neural Computation, 1(2):263–269.

Pedregosa, F., Varoquaux, G., Gramfort, A., Michel, V., Thirion, B., Grisel, O., Blondel, M., Prettenhofer, P., Weiss, R., Dubourg, V., Vanderplas, J., Passos, A., Cournapeau, D., Brucher, M., Perrot, M., and Duchesnay, E. (2011). Scikit-learn: Machine learning in Python. Journal of Machine Learning Research, 12:2825–2830.

Pessoa, L. and Ungerleider, L. G. (2004). Top-down mechanisms for working memory and attentional processes.

Scholkopf, B. and Smola, A. J. (2001). Learning with kernels: support vector machines, regularization, optimization, and beyond. MIT press.

Schurz, M., Radua, J., Aichhorn, M., Richlan, F., and Perner, J. (2014). Fractionating theory of mind: a meta-analysis of functional brain imaging studies. Neuroscience & Biobehavioral Reviews, 42:9–34.

Shine, J. M., Breakspear, M., Bell, P., Martens, K. E., Shine, R., Koyejo, O., Sporns, O., and Poldrack, R. (2018). The low dimensional dynamic and integrative core of cognition in the human brain. bioRxiv, page 266635.

Skowronski, M. D. and Harris, J. G. (2007). Automatic speech recognition using a predictive echo state network classifier. Neural networks, 20(3):414–423.

Smith, J. F., Hur, J., Kaplan, C. M., and Shackman, A. J. (2018). The impact of spatial normalization for functional magnetic resonance imaging data analyses revisited. bioRxiv, page 272302.

Steil, J. J. (2004). Backpropagation-decorrelation: online recurrent learning with o (n) complexity. In Neural Networks, 2004. Proceedings. 2004 IEEE International Joint Conference on, volume 2, pages 843–848. IEEE.

Sussillo, D. and Abbott, L. F. (2009). Generating coherent patterns of activity from chaotic neural networks. Neuron, 63(4):544–557.

Triefenbach, F., Jalalvand, A., Schrauwen, B., and Martens, J.-P. (2010). Phoneme recognition with large hierarchical reservoirs. In Advances in neural information processing systems, pages 2307–2315.

Vandoorne, K., Dierckx, W., Schrauwen, B., Verstraeten, D., Baets, R., Bienstman, P., and Van Campenhout, J. (2008). Toward optical signal processing using photonic reservoir computing. Optics express, 16(15):11182–11192.

Williams, R. J. and Zipser, D. (1989). A learning algorithm for continually running fully recurrent neural networks. Neural computation, 1(2):270–280.

Yu, B. M., Cunningham, J. P., Santhanam, G., Ryu, S. I., Shenoy, K. V., and Sahani, M. (2009). Gaussian-process factor analysis for low-dimensional single-trial analysis of neural population activity. In Koller, D., Schuurmans, D., Bengio, Y., and Bottou, L., editors, Advances in Neural Information Processing Systems 21, pages 1881–1888. Curran Associates, Inc.

Zacks, J. M., Braver, T. S., Sheridan, M. A., Donaldson, D. I., Snyder, A. Z., Ollinger, J. M., Buckner, R. L., and Raichle, M. E. (2001). Human brain activity time-locked to perceptual event boundaries. Nature neuroscience, 4(6):651.

